# Paired single-cell host profiling with multiplex-tagged bacterial mutants reveals intracellular virulence-immune networks

**DOI:** 10.1101/2022.03.06.483158

**Authors:** Ori Heyman, Dror Yehezkel, Neta Blumberger, Gili Rosenberg, Camilla Ciolli Mattioli, Aryeh Solomon, Dotan Hoffman, Noa Bossel Ben-Moshe, Roi Avraham

## Abstract

Encounters between host cells and intracellular bacterial pathogens lead to complex phenotypes that determine the outcome of infection. Single-cell RNA-sequencing (scRNA-seq) are increasingly used to study the host factors underlying diverse cellular phenotypes. But current approaches do not permit the simultaneous unbiased study of both host and bacterial factors during infection. Here, we developed scPAIR-seq, an approach to analyze both host and pathogen factors during infection by combining multiplex-tagged mutant bacterial library with scRNA-seq to identify mutant-specific changes in host transcriptomes. We applied scPAIR-seq to macrophages infected with a library of *Salmonella* Typhimurium secretion system effector mutants. We developed a pipeline to independently analyze redundancy between effectors and mutant-specific unique fingerprints, and mapped the global virulence network of each individual effector by its impact on host immune pathways. ScPAIR-seq is a powerful tool to untangle bacterial virulence strategies and their complex interplay with host defense strategies that drive infection outcome.

## Introduction

Interactions between a pathogenic intracellular bacterium and its host involve both activation of a coordinated defense response by the host macrophage and a complex virulence program executed by the bacterium (Schwan et al., 2000). To invade and survive within the hostile host cellular environment, intracellular pathogens employ secretion systems that deliver effector proteins across phospholipid membranes from within the bacterium into the host cytoplasm (Green and Mecsas, 2016). A wide range of secretion systems allow different pathogens to survive within host cells. For example, in *Mycobacterium tuberculosis*, a type VII secretion system (T7SS) is required for reprogramming of host cell death mechanisms (Stanley et al., 2003). In *Salmonella* Typhimurium (*S*.Tm), a type III secretion system (T3SS) can remodel host phagosomes into a hospitable vacuole, whereas the T3SS of *Shigella flexneri* enables bacteria to lyse the phagocytic membrane and gain access to the cytoplasm shortly after internalization (Hueck, 1998). *Legionella pneumophila* encodes a type IV secretion system that injects ∼300 effectors, allowing the pathogen to subvert host processes and establish replication-permissive niche inside an ER-like vacuole (Tilney et al., 2001).

One of the best studied intracellular pathogens, *S*.Tm utilizes two different T3SS, located on Salmonella Pathogenicity Island (SPI) 1 and 2, to establish a protective niche within different host cell types (Hansen-Wester and Hensel, 2001). SPI-2 mediates secretion of 28 effectors from within the macrophage phagosome to manipulate host cell processes and ensure bacterial replication inside the *Salmonella*-containing vacuole (SCV) (Jennings et al., 2017). In recent years it has become increasingly understood that effector secretion is heterogeneous between individual intracellular bacteria, leading to different complex phenotypes (García-del Portillo and Pucciarelli, 2017). For example, within a population of invading *S*.Tm, variable activity of PhoP, a response regulator required for SPI-2 T3SS activation (Bijlsma and Groisman, 2005), has been shown to drive heterogeneity in macrophage responses (Avraham et al., 2015). Heterogeneity was detected within *S*.Tm subpopulations comprising either actively growing bacteria or non-growing persisters, with different SPI-2 expressing profiles in infected host cells (Stapels et al., 2018). Within a single population, both in vitro and in vivo, *S*.Tm was shown to display heterogeneous *S*.Tm T3SS expression has been associated with variable growth rate (Sturm et al., 2011), sensitivity to antibiotics (Arnoldini et al., 2014) and localization of bacterial cells within infection sites (Laughlin et al., 2014). Adding to the complexity of heterogenous expression of T3SS and the host response, redundancy between effectors activities is also observed, as single T3SS effectors converge on the same host pathway (Galán, 2009). For example, the GTPase-activating protein SopD2 and the cysteine protease GtgE were both shown to target and counteract the host Rab32-dependant host defenses (Wachtel et al., 2018). Recent studies have additionally shown that a single *S*.Tm effector can bind many host proteins that are associated with multiple cellular processes (Walch et al., 2021). While redundancy and heterogeneity are beneficial for the pathogen to establish a successful infection, they have complicated the determination of individual effector function; therefore, a systematic understanding of T3SS effector activity is still lacking.

On the host side, macrophages infected with *S*.Tm generates well-documented heterogeneous phenotypes during intracellular infection (Burton et al., 2014). Recent single cell RNA-seq (scRNA-seq) studies revealed host signatures of macrophage subpopulations that display diverse outcomes: some are permissive to intracellular bacterial growth (Hoffman et al., 2021); some will lyse the ingested bacteria (Huang et al., 2018); others allow bacteria to persist intracellularly (Stapels et al., 2018). But scRNA-seq methods are limited to profiling only host transcripts as they rely on polyA priming of mRNAs, while bacterial mRNAs lack polyA. The heterogeneous and dynamic nature of bacterial pathogens suggests that descriptions limited only to immune attributes may fail to accurately characterize the full spectrum of interactions with different complex phenotypes (Patel et al., 2021). Recent efforts to perform single cell profiling of bacteria have had limited sensitivity for detection of bacterial genes, making biological interpretation in the context of infection difficult (Avital et al., 2017; Imdahl et al., 2020; Kuchina et al., 2021). Alternatively, systematic analysis of bacterial gene function can be performed *en masse* either as arrayed or pooled genetic screens, but are mostly limited to autonomous phenotypes rather than interactions with host and are insensitive to bacterial heterogeneity (Barczak et al., 2017; O’Connor and Isberg, 2014). Thus, a significant gap remains in our ability to study macrophage and bacterial factors that regulate survival strategies during individual encounters. Knowledge of these factors is fundamental to our understanding of infection biology and finding novel treatment options for infectious disease that result in a more beneficial outcome to the host.

To address this challenge, we introduce scPAIR-seq, a computational and experimental single cell framework for analyzing PAthogen-specific Immune Responses. This approach offers the much-needed simultaneous, systematic analysis of host and bacterial factors that contribute to heterogeneous host-pathogen encounters. We implemented a unique multiplexed bacterial Mutant Identifier Barcode (MIB) and modified scRNA-seq protocols for detection of both the bacterial mutant identity and the transcriptional profile of single infected host cells (**Fig. 1**). As a proof-of-principle, we generated a MIB-tagged library of *S*.Tm SPI-2 effector mutants and followed their cognate effects on the host using the transcriptome of single infected macrophages. We developed a computational pipeline to analyze mutant-dependent changes in host activation programs and identified specific host targets, gene signatures, and cell states affected by individual SPI-2 mutants. Using this approach, we show that a SPI-2 effector, SifA, has a unique, yet undescribed function, that drive a distinct macrophage transcriptional signature.

**Figure 1:**
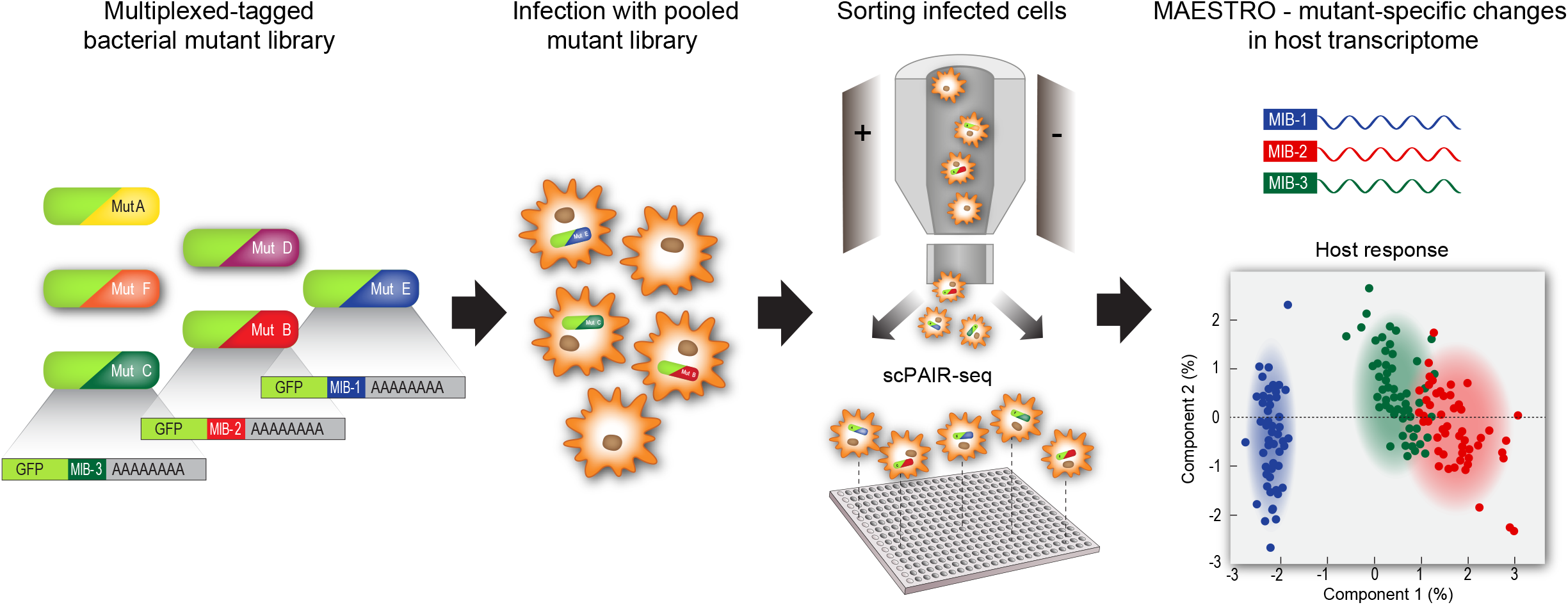
Overview of the scPAIR-seq approach. Bacterial mutants express a fluorescent protein (GFP), tagged with unique mutant identifier barcode (MIB) followed by a polyA sequence. Macrophages were infected with a library of pooled multiplex-tagged bacterial mutants and single cells were sorted into multi-well plates. scPAIR-seq was applied on infected cells, for deconvolution of mutant-specific changes in host transcriptome using the MAESTRO pipeline.

## Results

### Establishing a scRNA-seq protocol for detection of MIBs in intracellular bacteria

To allow detection of bacterial mutants within single host cells, we engineered a plasmid with constitutive expression of a Green Fluorescent Protein (GFP), tagged with a MIB followed by a polyA tail, and inserted it into *S*.Tm (MIB-*S*.Tm; **Fig. 1**). The GFP signal enables detection of infected cells using flow cytometry and imaging. The polyA tail allows capture of MIBs using scRNA-seq protocols, and deconvolution of bacterial mutant identity by the unique bacterial MIB transcripts. Polyadenylation in bacteria is considered a signal for transcript degradation by a mechanism involving 3’-exonucleolytic attack (Hui et al., 2014). To assess the impact of the polyA tag on transcript abundance, we compared GFP fluorescence levels between MIB-*S*.Tm to *S*.Tm containing non-polyadenylated GFP plasmid (*S*.Tm control), and *S*.Tm without GFP plasmid (No-GFP control; **fig. S1A**). While there is a decrease in the GFP fluorescence of MIB-*S*.Tm compared to *S*.Tm control, GFP signal remains sufficiently high in this strain for detection within host cells. We next tested if the GFP with MIB transcript can be detected by polyA capture of bacterial RNA. We detected GFP with MIB transcript only in the MIB-*S*.Tm samples but not in *S*.Tm control, validating that polyadenylated MIB transcripts were amplified exclusively (**fig. S1B**).

Next, as bacterial transcripts within single infected host cell are estimated at less than 1% of the total RNA (Marsh et al., 2017), we sought to modify a scRNA-seq protocol (Hashimshony et al., 2016) to specifically amplify low abundant MIB transcripts (**Fig. 2A**). After an in vitro transcription (IVT) step, the sample is split into two parts and amplified by PCR separately to generate both i) a library of host transcripts via traditional protocol, and ii) a MIB library with specific amplification of the tagged GFP. The MIB library is then sequenced with a custom primer that targets the GFP transcript upstream of the MIB sequence to allow exclusive detection of MIBs. The adjusted protocol was then tested for sensitivity and accuracy of host transcriptome analysis and MIB identification in bulk. We infected a J774.1 cells with MIB-*S*.Tm, collected 150 cells by fluorescence activated cell sorting (FACS) and generated bacterial MIB libraries. To test the sensitivity of MIB detection, we analyzed MIB reads and counted ∼625,000 reads of ∼1,400,000 reads in the MIB-*S*.Tm bulk sample that was generated from 150 cells. Second, we generated MIB libraries from serial dilutions of RNA samples and measured detection of MIBs within host cells (**fig. S2A**).

**Figure 2:**
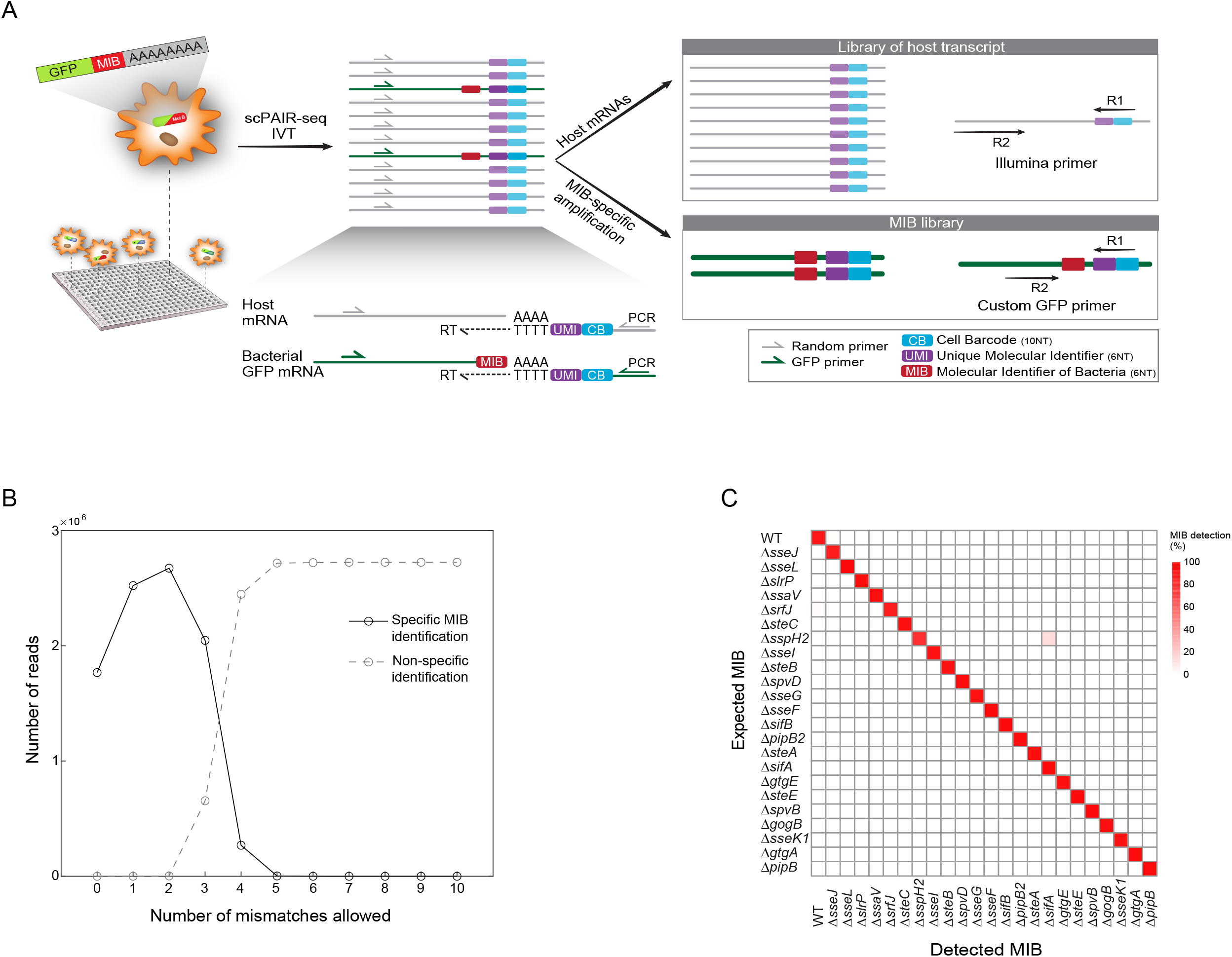
scPAIR-seq protocol and optimization for paired analysis of MIBs and host transcripts. **(A)** Detailed overview of scPAIR-seq protocol. UMI, unique molecular identifier; CB, cell barcode; RT, reverse transcription; NT, nucleotides **(B)** Cultures of multiplexed-tagged SPI-2 mutants were pooled and MIB library was generated. Sequences from each read were compared to the MIB sequence of each mutant with increasing number of mismatches allowed from 0 to 10. Indicated is the number of specific identifications (each read mapped exactly to one mutant; solid black line) and number of non-specific identifications (more than 1 mutant mapped to the same read; dashed gray line). Using one mismatch we optimize MIB identification rate without affecting specificity. **(C)** BMDMs were infected separately with each of the *S*.Tm SPI-2 mutants and bulk samples were collected. Percentage of the bacterial MIBs detected (columns) are presented for each infected sample (rows); see colorbar to the right. Each detected MIB was normalized by the total number of reads obtained for that MIB across all infected samples, before calculating percentages.

### Construction of a pooled, MIB-tagged, SPI-2 mutant library

We next aimed to test scPAIR-seq by performing functional analysis of SPI-2 effectors with single cell resolution. We generated and validated a library of WT and 24 known SPI-2 effector mutants (Porwollik et al., 2014) and transformed each mutant with a GFP expressing plasmid containing a unique MIB sequence (**table S1**). To minimize PCR and sequencing errors that can lead to MIB misidentification (Sleep et al., 2013), we designed MIB sequences with an edit distance of at least three nucleotides (**fig. S2B**), providing two mismatches for non-ambiguous identification of mutants. To examine MIB classification and detection, we pooled cultures of bacterial mutants, extracted RNA and generated MIB libraries. To test for specificity of MIB detection, we iteratively increased number of mismatches that are allowed for MIB classification (**Fig. 2B and fig. S2C**). We could validate specific MIBs classification when allowing up to 2 mismatches, corresponding to our design of MIBs. Allowing more than 2 mismatches resulted in ambiguous MIB detection. We next verified the specificity and sensitivity of detection of bacterial MIBs in infected bone marrow-derived macrophages (BMDMs). We infected BMDMs with each of SPI-2 mutants separately, sorted 150 infected cells from each sample and generated MIB libraries. We measured highly specific detection of the expected MIBs from each infected bulk sample, according to its infected mutant (**Fig. 2C**). Furthermore, high signal to noise ratio was observed for each individual MIB (**fig. S2D**). MIB of ΔSopD2 was not detected and was verified separately. As a control, a BMDM sample infected with *S*.Tm strain carrying a non-polyadenylated GFP plasmid showed no detection of MIBs.

### A computational framework for MIB identification within single cells

We next performed scPAIR-seq protocol on single BMDMs infected with the pooled library of SPI-2 effectors. We infected BMDMs with the pooled MIB-tagged SPI-2 mutant library at a low multiplicity of infection (MOI) of 2:1 to ensure an infection rate of ∼1 bacterium per cell (Gog et al., 2012). SPI-2 gene expression is induced approximately 1.5 hours post internalization (Jennings et al., 2017), peaks at 2-4 hours after infection (Fass and Groisman, 2009), and mediates bacterial replication after 18 hours (Nix et al., 2007). Thus, we harvested cells at twenty hours post infection (hpi) and isolated 3648 infected single-cells to deconvolute the phenotype of host responses corresponding to individual SPI-2 mutants. We index-sorted infected cells based on the GFP signal of the internalized bacteria. Bulk samples of 1000 infected cells were collected as controls for validating single-cell MIB detection. MIB and host libraries were generated and sequenced according to the scPAIR-seq protocol. For each single cell we extracted the corresponding MIB reads and generated a matrix of MIB counts per cell. Examining total MIB counts revealed a distribution with three distinct peaks (**fig. S3A**). We set a threshold at the maximum of the lowest peak to exclude cells with less than 128 detected MIB reads, reasoning that counts below this threshold would reflect low-quality detection. Cells with low-quality MIB detection were removed, and downstream analysis was performed on 1191 cells (∼33%). We developed a computational framework to determine unambiguous mutant identity in single cells with scPAIR-seq. We modeled the normalized counts of each MIB to find the quantile parameter that maximizes singlet detection rate across all cells in the experiment. We applied ‘deMULTIplex’ (McGinnis et al., 2019) to the MIB counts within single host cells and set a quantile threshold of 0.23 that maximizes the percentage of cells containing a single mutant MIB (**Fig. 3A**). Importantly, this threshold is robust to singlet detection at a range of threshold values. Based on this analysis, we assigned 878 cells with single invading mutant and 112 cells with more than one mutant (**Fig. 3B**).

**Figure 3:**
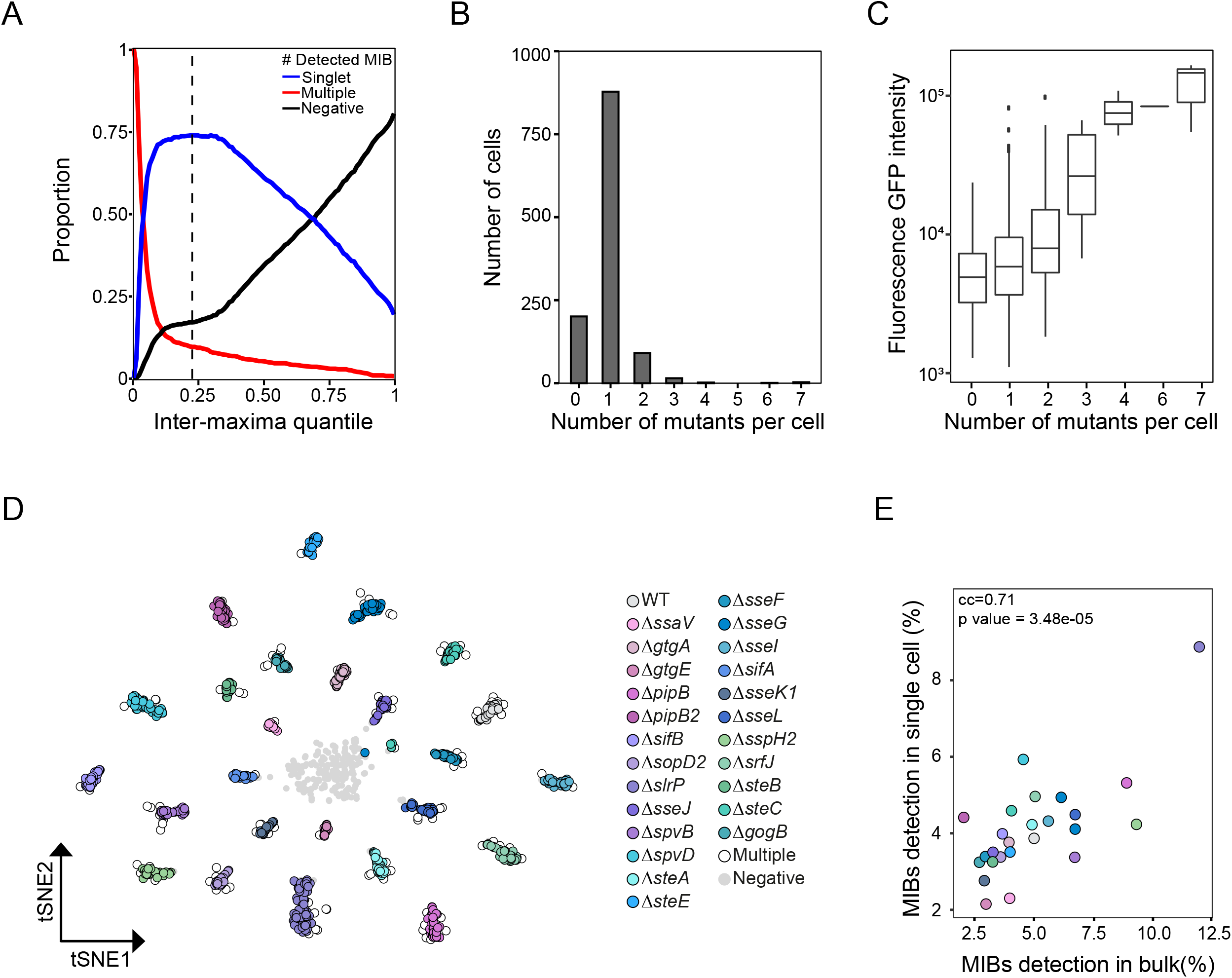
Non-ambiguous identification of MIBs in single infected cells. BMDMs were infected with a pooled MIB-tagged SPI-2 mutant library at an MOI=2. Twenty hpi single infected host cells were sorted and analyzed by scPAIR-seq. (**A**) The distribution of MIB counts across all cells was modeled and the local maxima of positive and negative cells were found for each MIB (see supplementary methods). To set a threshold between these two maxima of each MIB, in a way which maximizes singlets detection across all MIBs together, we iterated over all inter-maxima quantiles (x-axis). The percentage of singlets (infected cells with 1 mutant; blue line), multiple (more than 1 invading mutant per cell; red line), and negative (no invading mutant was identified; black line) are shown (y-axis) for each quantile. We selected the quantile which optimize singlet detection (q=0.23, black dashed line). (**B**) Number of detected mutants per cell indicate that most cells are infected with 1 mutant as expected by the low MOI. (**C**) Sorted single cells were indexed by the GFP signal of internalized bacteria. Boxplots represent the GFP intensities (log_10_) from host cells as detected by flow cytometry (y-axis), grouped by the number of MIBs detected by scPAIR-seq in single cells (x-axis). There is high concordance between the number of identified invading bacteria and the GFP intensity. (**D**) Visualization of MIB counts in single cells using t-distributed Stochastic Neighbor Embedding (t-SNE) dimensional reduction. Each cell is colored by the identity of its invading mutant as detected by scPAIR-seq (see colorbar to the right and fig. S3C). Depicted are also cells with multiple MIB detection (white) or without detection (Negative; gray). (**E**) Comparison between MIB detection from the single cell data (y-axis) and detection in matched bulk infected samples (x-axis; averaged distribution from 4 samples). Dots represent the percentage of each mutant from total MIBs. Colors are as in (D), correlation coefficient and corresponding p-value are indicated in the figure.

Few cells were detected with 4-7 invading mutants, raising the possibility that these cells were simultaneously invaded by multiple mutants during early stages of infection. Alternatively, infection with multiple mutants could arise from re-infection; while gentamycin-containing media was used throughout the infection course, re-infection events can occasionally occur (Stapels et al., 2018). As another alternative, multiple infecting mutants could result from efferocytosis, a process in which a macrophage swallows another dying infected cell (Korns et al., 2011). To validate multiple MIBs detection in single cells, we assessed the indexed GFP intensities of the respective sorted single cells. Indeed, GFP intensity levels were higher in cells with higher number of MIBs (**Fig. 3C**). To further validate that the GFP signal serves as a proxy for bacterial numbers, we analyzed infected cells and measured increased GFP fluorescence at twenty hours compared to four hours **(fig. S3B**), by which time bacterial replication has occured. Thus, using GFP fluorescence within infected cells we can evaluate bacterial burden and identify single cells that are infected by multiple mutants.

Next, we generated a map to capture the landscape of MIBs in single infected host cells. t-distributed Stochastic Neighbor Embedding (tSNE) dimensional reduction was used on the MIB counts to represent the detection in the single cell data (**Fig. 3D**). This dimensional reduction allows visualization of the cells that were infected by each mutant, based on our pipeline for MIB identification. Overall, we identified between 19 to 76 infected cells for each mutant (**fig. S3C**). To validate our procedure for MIB identification in single cells, we quantified MIBs from matched bulk infected samples. We averaged the distribution of detected MIBs from 4 bulk samples and compared it to MIB detection in the single cell data. We found a high correlation (correlation coefficient of 0.71), corroborating our single cell detection (**Fig. 3E**). To further confirm MIB library analysis, we analyzed the detection of MIBs also in host libraries. We found a high degree of agreement in the detection of MIB identity between host libraries and MIB libraries. The number of MIB-detected cells was ∼5-folds higher in the MIB libraries (**fig. S3D**), confirming the improved sensitivity of our protocol. Taken together, we demonstrate an approach to detect unique MIBs that map bacterial mutant identity within single host cells.

### A computational framework to dissect mutant-specific changes of host transcriptome

To study distinct host cell responses affected by individual SPI-2 effector mutants, we next analyzed host transcriptomes in single cells. We first validated quality control parameters of host single cell data (**fig. S4; A-C**), and turned to analysis of single cells. Unsupervised clustering analysis showed that host gene-expression profiles formed six stable clusters (**fig. S4D**). To study the possibility of mutant-specific clusters, we examined the composition of the clusters across cells that were grouped based on the identity of the infected mutant (**fig. S4E**). Macrophages infected with WT *S*.Tm express three main clusters of genes: cluster Ⅰ containing inflammation-related genes (e.g., *Il1b, Nfkbia* and *Cxcl2*), cluster Ⅲ containing genes related to cell cycle (e.g., *Cdk1, Mcm6* and *Lig1*) and cluster Ⅴ with genes related to oxidative stress (e.g., *Prdx1, Prdx6* and *Nqo1*). These clusters were also shared with cells infected by most mutants, indicating the known redundancy between effectors (Zhou et al., 2001). Three clusters (Ⅱ, Ⅳ and Ⅵ) were mostly evident in several mutant-infected cells. Of note, clusters Ⅱ and Ⅵ were most evident in Δ*steE* and Δ*sifA* infected cells, respectively. Cluster Ⅱ, which was expressed mainly by Δ*steE* infected cells, includes the type Ⅰ interferon (IFN) response genes (e.g., *Oasl2, Ifitm6* and *Cd74*) and cluster Ⅵ, associated mainly with Δ*sifA*, was enriched with genes reminiscent of M2-transcriptional state (e.g., *Il4ra, Arg1* and *Mrc1*) (Saliba et al., 2016). We next performed a directed analysis of the impact of individual SPI-2 mutants on host gene expression. To identify changes due to mutant identity, we developed a tailored pipeline for single cell analysis using a metric we termed MutAnt spEcific hoST TRanscriptOme (MAESTRO) analysis. The analysis estimates the effect of each mutant on host gene expression changes by weighing both expression levels and variance of each gene between WT-and mutant-infected cells. We first modeled the number of cells that are required to represent the variance of the data, to ensure that the cell numbers in each mutant group is sufficient. In agreement with previous estimations (Dixit et al., 2016), we observed that from 20 cells the variance across cells is stabilized **(fig. S4F)**. Next, infected cells were grouped by MIB identity (25 groups) and mean expression and coefficient of variance (CV) were calculated for all genes in each mutant group. To generate a metric that captures these two measurements of single cell data, we calculated the Euclidean distance in mean-CV space between WT and respective mutant (MAESTRO score) for each gene (**Fig. 4A**). To exclude noise due to lowly detected genes, we included only genes expressed in at least 30% of cells for specific mutant. As lowly expressed genes in scRNA-seq tend to display high CV due to dropouts, we controlled artificial high CV by increasing the weight for mean expression in the analysis. Plotting the MAESTRO scores distribution for all expressed genes in all mutants, we observed a right tail of genes with high MAESTRO scores (**Fig. 4B**). We curated gene lists linking each bacterial mutant with its effect on host gene expression. We defined a group of Differentially Expressed Genes (DEGs) for each mutant by selecting the genes with MAESTRO scores higher than the 95^th^ percentile of all scores (**fig. S5** and **table S2**).

**Figure 4:**
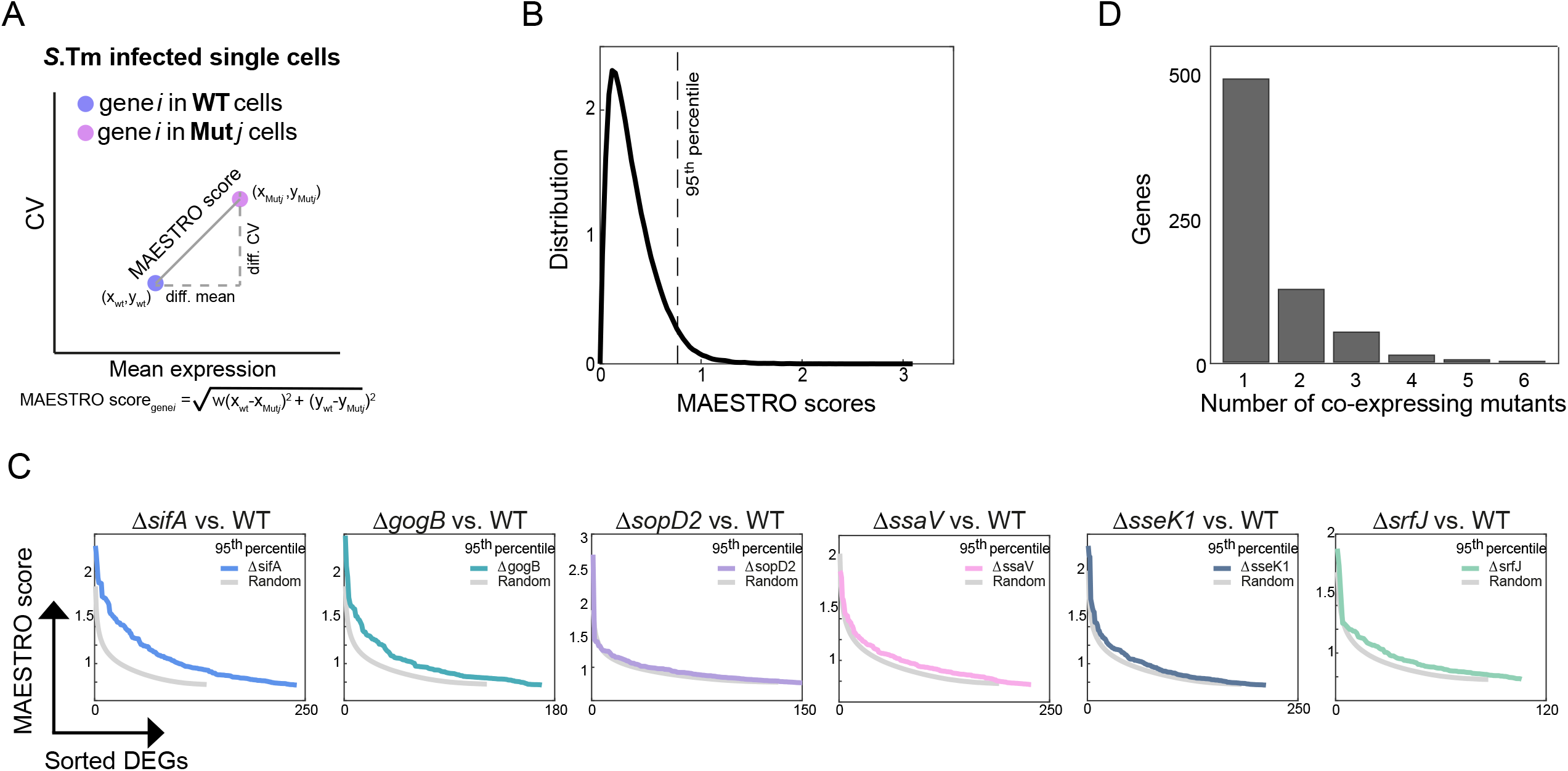
MAESTRO analysis to dissect mutant-specific changes of host transcriptome. **(A)** Illustration of the MAESTRO scores calculation: to capture mutant-specific changes in host gene expression, we calculated the mean expression (x-axis) and CV (y-axis) of each gene (*i*) in each mutant group (*j*) compared to WT infected cells. The MAESTRO score was calculated as a weighted Euclidian distance between the mean-CV of the WT and the specific tested mutant (*j*), with weight (*w*) for the mean expression component set to 1.5. (**B**) Histogram of MAESTRO scores from all genes across all mutant groups generated a right tail distribution of genes with high MAESTRO scores. We defined the genes with MAESTRO score larger than the 95^th^ percentile of all scores (black dashed line) as different from WT infected cells. Using this threshold, we defined the group of differentially expressed genes (DEG) for each mutant. (**C**) To test for significance of these DEGs we generated a random model. We compared the distribution of MAESTRO scores obtained from our data to the null distribution. Using two-sample t-test we identified 6 mutants with significantly higher MAESTRO scores than the random model. Presented are the scores from these 6 significant mutants (colored lines), and the random scores of each mutant (gray lines; mean of 1000 random permutations) (**D**) To identify specific and shared signatures, DEGs of the six significant mutants were intersected. Bar plots represent the number of genes that are unique to one mutant, and these that are shared between 2 to 6 mutants (x-axis).

To provide statistical power for MAESTRO scores and test for mutants that significantly impact host genes relative to WT, we generated a random model. We included in our random model cells that were excluded from the host analysis due to low MIB coverage (**fig. S3A**, n=2457). We randomly assigned these cells to groups in the same sizes as the original number of cells we obtained for each mutant. We calculated MAESTRO scores for these random mutant groups and generated the null distribution of the scores. We then performed two-sample t-test between our real data and the random data for each mutant. Using this strategy, we identified 6 mutants with significantly different MAESTRO scores than the random model (1% FDR), including Δ*sifA*, Δ*gogB*, Δ*sopD2*, Δ*ssaV*, Δ*sseK1* and Δ*srfJ* **(Fig. 4C)**. We identified distinct signatures of each of these mutants, with 495 out of total 691 DEGs (∼72%) that were mutant specific (**Fig. 4D, table S3**). MAESTRO, our unique computational framework, thus enabled us to analyze both redundancy between mutants and to assign individual mutant-specific host gene expression fingerprints.

### Functional analysis of effectors by mutant-specific host gene expression fingerprints

We performed Gene Ontology (GO) enrichment analysis on the groups of DEGs for each significant mutant. A total of 24 terms were enriched (10% FDR), with 15 terms that were mutant specific (**Fig. 5A, table S4**). This analysis identified to previously described functions of these effectors, providing validation of our approach. General inflammatory response GO-terms (enriched in Δ*sifA*, Δ*sseK1* and Δ*gogB*) and neutrophil chemotaxis (enriched in Δ*sifA*, Δ*sseK1*, Δ*gogB* and Δ*sopD2*) indicated a role for these effectors in modulating innate immune response, as previously described (Jennings et al., 2017). Cellular response to tumor necrosis factor (TNF), a well-characterized inducer of NF-kB (Hayden and Ghosh, 2014), was enriched only in Δ*sseK1* and Δ*gogB*. These two effectors were previously identified as targeting the NF-kB signaling pathway. SseK1 was shown to suppress TNFα-induced NF-kB activation through direct modulation of the host signaling adaptor TRADD (Günster et al., 2017; Newson et al., 2019), and GogB was reported to down-regulate host inflammatory responses by inhibiting poly-ubiquitination of IkBα and thus NFkB-dependent gene expression (Pilar et al., 2012). The effector SifA is known to maintain the integrity of the SCV membrane (Beuzón et al., 2000), which is indicated in our analysis by signatures of Δ*sifA* related to endocytosis, a necessary process for SCV maturation (Smith et al., 2005). In addition, response to oxidative stress was observed in our analysis of Δ*sifA*, corresponding to increased redox stress described in Δ*sifA* infected cells (Heijden et al., 2015). SopD2 has been shown to interact with various host regulatory Rab-GTPases in the endocytic pathway and block its function (D’Costa et al., 2015; Spanò et al., 2016; Teo et al., 2017). Rab GTPases mediate the motility of organelles and vesicles and contribute to membrane trafficking (Gillingham et al., 2014). In our analysis we identified enrichment of Δ*sopD2* with ER to Golgi vesicle-mediated transport. We next calculated the mean expression of up or down regulated DEGs of each mutant across all cells (**Fig. 5B**). We measured high variation within each group of cells infected with the same mutant, indicating signature expression in only a subpopulation of cells (**Fig. 5C**). T3SS effectors are considered to form a robust, redundant intracellular signaling network that could sustain deletion of a single effector gene (Ruano-Gallego et al., 2021). Thus, the observed heterogeneity in host response possibly reflects this redundancy in effectors network, which can mask the deleted effector activity in a subset of infected cells. In cells infected with Δ*ssaV*, a mutant with nonfunctional SPI-2, we measured reduced variation in host responses, potentially reflecting ablated redundancy of the entire effector network (**Fig. 5C**). Overall, we provide a gene signature analysis with mutant-specific fingerprints driving unique changes in host gene expression.

**Figure 5:**
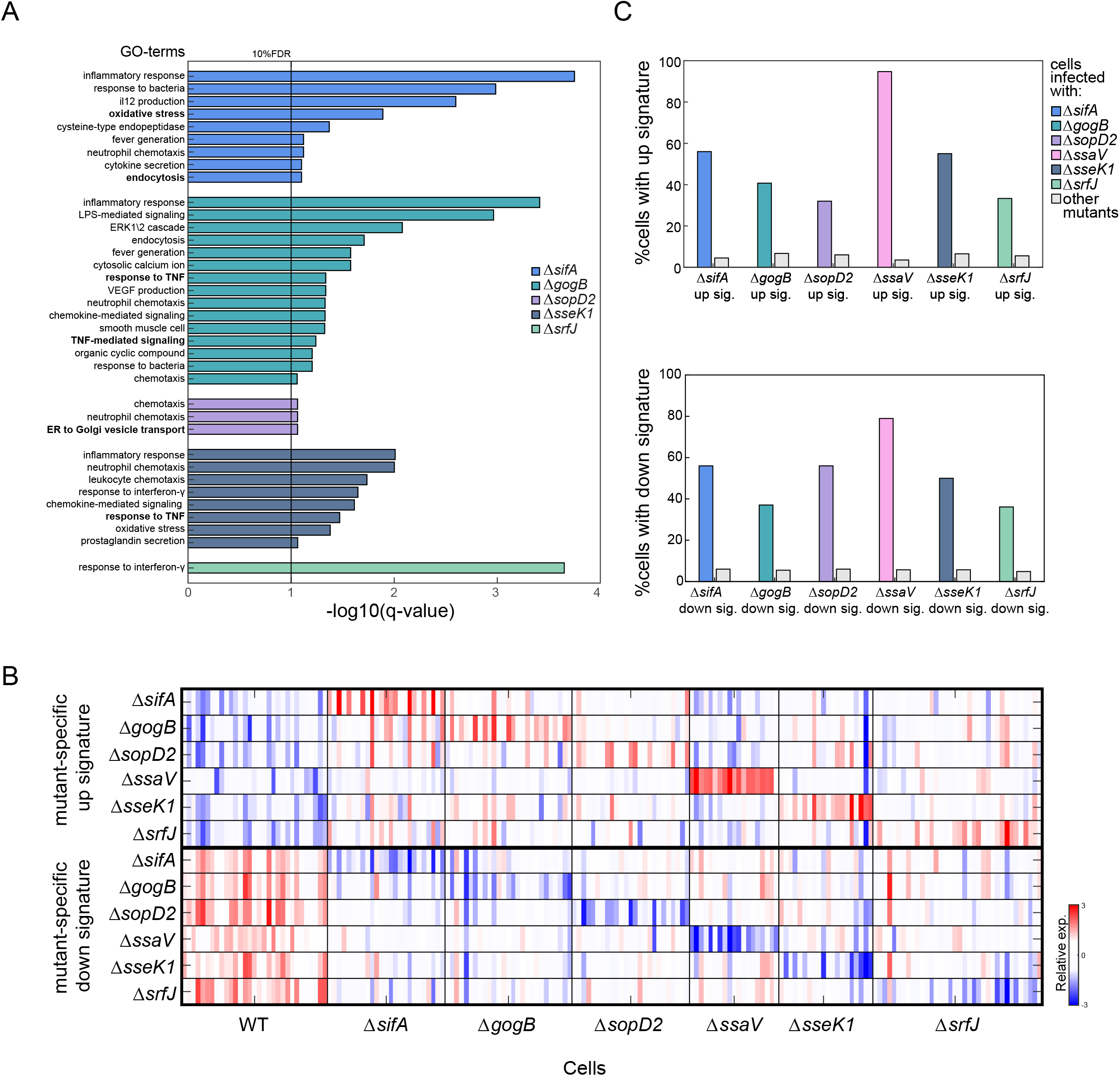
MAESTRO analysis reveals virulence-immune networks of SPI-2 mutants. **(A)** GO-terms enrichment analysis for the DEGs of each significant mutant (**table S4**) **(B)** Heatmap of the mean expression of mutant-specific signature genes (up – top panel, down – bottom panel regulated DEGs) for each mutant across all cells. Rows are the mean expression of the up or down regulated genes of each mutant (mutant signature); columns are sorted by the MIB detected in infected single cells. **(C)** For each up or down regulated signature of each mutant, we calculated the number of cells expressing the signature. Presented are the proportion of cells that up-regulated (upper panel) or down-regulated (bottom) the specific signature in the same mutant (colored bars), or in all other cells not infected with this mutant (gray bars, non-specific induction of the signature). Signatures are specific to their mutant infected cells, with high variation between single cells.

### SifA effector protein drives an alternatively activated macrophage phenotype

We next interrogated selected effector function using its host transcriptional fingerprints. Based on MAESTRO analysis and clustering of host transcriptome, the SifA mutant presented the most prominent phenotype. SifA is a well-studied SPI-2 effector, known to play a significant role in *S*.Tm virulence: detoxifying lysosomes (McGourty et al., 2012), recruiting late endosomes and lysosomes to the SCV (McEwan et al., 2015), and promoting the formation of tubular membranous structures connected to SCVs (Portillo et al., 1993; Stein et al., 1996). These studies, like for other T3SS effectors, focus on characterizing the host protein targeted by SifA. Interestingly, when examining the transcriptional differences between WT and Δ*sifA* within infected cells (**Fig. 6A**), we found enrichment in genes related to macrophages polarization (M1/M2 genes, **table S5**, taken from Saliba et al., 2016), corresponding to cluster Ⅵ in the unsupervised single cell analysis (**fig. S4; D and E**). Furthermore, M2 host genes are exclusively expressed by only a subset of Δ*sifA* infected cells (**Fig. 6B**).

**Figure 6:**
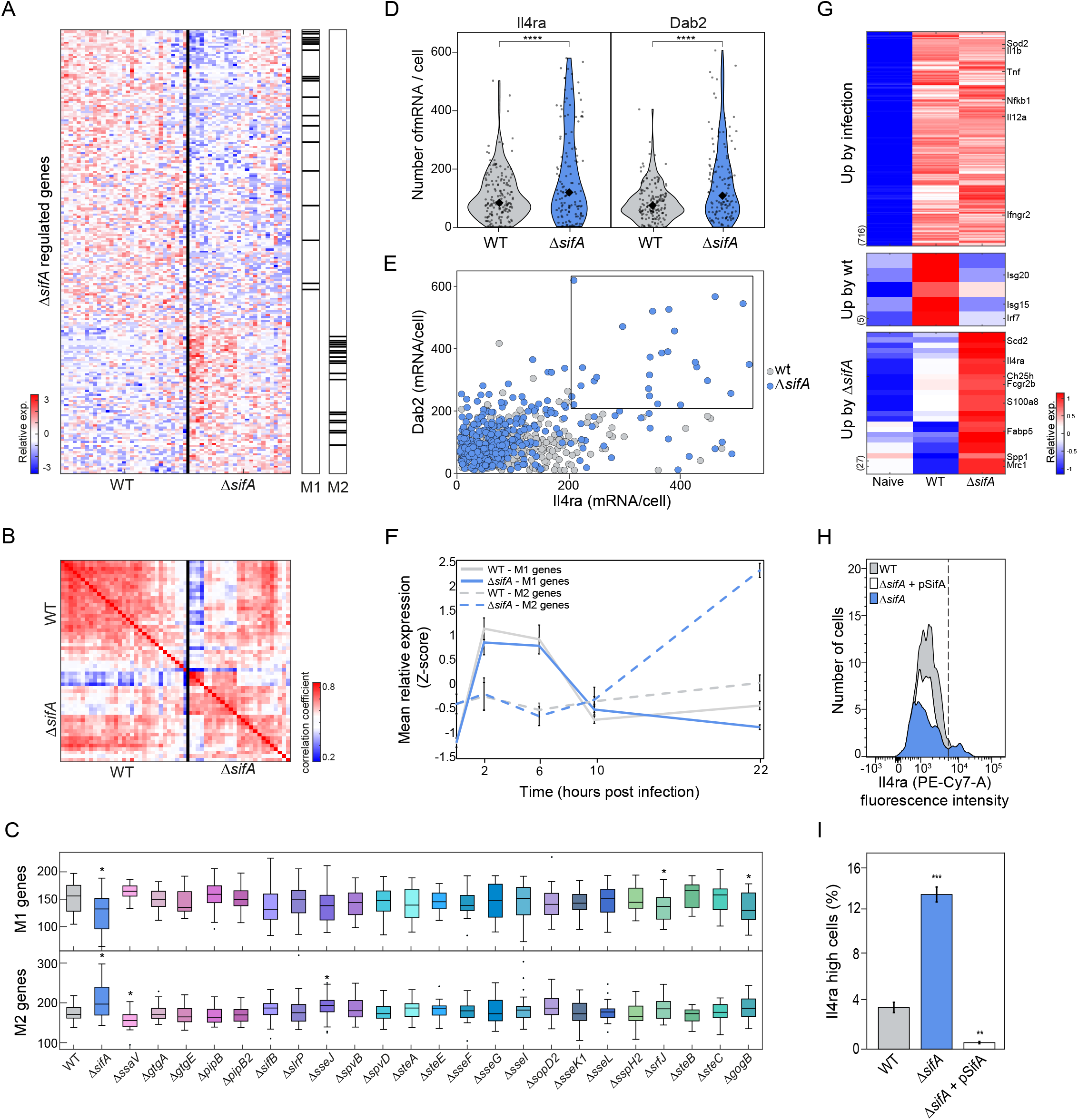
Δ*sifA* mediates transcriptional reprogramming of BMDMs towards M2-activation state. **(A)** Heatmap of the expression levels of Δ*sifA* DEGs in single cells across WT and Δ*sifA* infected BMDMs. Genes and cells are clustered by hierarchical clustering (ward linkage method); see colorbar to the left for expression levels. The black bars to the right indicate position of genes related to macrophages M1 and M2 polarization (from Saliba et al., 2016, see **table S5**). M1 signature is enriched in down-regulated genes in Δ*sifA* infected cells (P=2.8×10^−13^), while M2 signature is enriched in the up-regulated genes (P<1×10^−20^). **(B)** Correlation matrix between WT and Δ*sifA* infected cells over the space of Δ*sifA* DEG; colorbar to the right indicates correlation coefficient values. There are two subsets of Δ*sifA* infected cells, one that has similar M1 signature to WT infected cells and another subset which elevates M2 polarization genes. **(C)** Box plots of the sum expression of M1 (upper panel) or M2 (bottom panel) genes across WT or *S*.Tm SPI-2 mutants. The box represents the median and 25-75^th^ percentile, whiskers encompass all data points. Mutants which significantly differentiate M1 or M2 expression levels relative to WT infected cells are indicated (two-sample t-test; 10%FDR). **(D** and **E)** BMDMs were infected with WT (gray) or Δ*sifA* (blue) and cells were fixed and analyzed by smFISH with probes against *Il4ra* and *Dab2* transcripts. **(D)** Violin plots showing *Il4ra* (left) and *Dab2* (right) mRNA counts per cell for each infected sample. Each dot corresponds to a cell, the median is indicated by the black diamond. **** P<0.0001, two-sample t-test. **(E)** Expression levels of *Il4ra* and *Dab2* transcripts across WT and Δ*sifA* infected cells. Rectangle marks a subpopulation of Δ*sifA*-infected cells with high expression of both transcripts. **(F)** Z-score normalized expression of representative M1 (filled lines) and M2 (dashed lines) genes (see Fig. S6; D and E) from WT (gray) or Δ*sifA* (blue) infected samples at 2, 6, 10 and 22 hpi and before infection at t=0 (naïve) by qRT-PCR. Expression was normalized to *Rps*13 gene. **(G)** Bulk RNA-seq of sorted naïve, WT or Δ*sifA* infected BMDMs twenty hpi. Presented is a heatmap of genes up-regulated following infection with WT or Δ*sifA* mutant relative to naïve cells (10% FDR). Genes were classified into three classes based on their expression changes between the conditions. See colorbar to the right for relative gene expression. For each class indicated representative genes; upper – inflammatory genes, middle - Type Ⅰ IFN genes, bottom – M2 and cholesterol-metabolism genes. **(H** and **I)** Infected BMDMs samples were stained with Il4ra antibody. Expression levels were analyzed by FACS (**H**, dashed line represent threshold for a population with high Il4ra expression) and proportion of cells expressing high Il4ra levels were compared between infected samples (**I**, cells right of the dashed line in **H**).

In response to the presence of microbial products, macrophage activation is considered to polarize in two distinct programs: the pro-inflammatory M1 macrophages, and alternatively activated M2 macrophages with tissue remodeling and immune resolution functions (Lee, 2019). Recent studies have shown that a SPI-2 effector of *S*.Tm function to support macrophages M2 activation (Stapels et al., 2018) and that intracellular growth was associated with M2 macrophages (Saliba et al., 2016). To test the specificity of Δ*sifA*-mediated M2 signature in macrophage polarization, we measured the sum expression of M1 and M2 genes in infected single cells across all groups of mutants (**Fig. 6C**). Gene expression in Δ*sifA* infected cells was significantly lower in M1 genes and higher in M2 genes compared to WT. In Δ*ssaV* infected cells we measured an opposite trend with significantly lower M2 gene expression, which is in agreement with previous reports indicating that a Δ*ssaV* mutant is less capable of triggering M2 polarization (Stapels et al., 2018). To validate M2 genes in only a subset of Δ*sifA* infected macrophages, we used single-molecule fluorescence in situ hybridization (smFISH). We designed probes targeting two transcripts, Il4ra and Dab2. Both are known markers of M2 polarization (Adamson et al., 2016) and both were upregulated in the Δ*sifA* signature (**table S2**). Similar to the scPAIR-seq results, we measured significantly higher *Il4ra* and *Dab2* expression in a subpopulation of Δ*sifA* infected cells compared to WT (**Fig. 6; D and E**, and **fig. S6A**). Expression levels of normalizing control gene *Gapdh* were comparable between WT and Δ*sifA* cells (**fig. S6B**). Moreover, when counting the number of bacteria within each infected cell, macrophages with higher loads of Δ*sifA* bacteria, but not WT, showed increased expression of *Il4ra* and *Dab2* (**fig. S6C**), indicating that the observed M2 gene program is not merely related to bacterial burden but rather related to Δ*sifA*-dependent activation.

Next, we characterized the kinetics of M1/M2 activation in Δ*sifA* infected macrophages. We studied the expression of four representative M1 genes (*Tnf, Cd40, Cxcl10* and *Nlrp3*) and measured early induction from 2 to 6 hpi with subsequent decrease in expression, both in WT and Δ*sifA* (**Fig. 6F** and **fig. S6D**). We then studied the expression of four representative M2 genes (*Il4ra, Dab2, Mrc1* and *Timp1*) and measured increased expression of all M2 genes in Δ*sifA* infected cells compared to WT infected cells 22 hpi (**Fig. 6F** and **fig. S6E**), indicating that M2 activation is triggered by Δ*sifA* only at later time points after infection. Next, we sorted bulk samples of WT or Δ*sifA* infected cells and performed bulk RNA-seq analysis. M2-related genes were significantly enriched in the genes that are up-regulated in Δ*sifA* relative to WT infected cells (P=2.2204e-15; ∼40% of genes) (**Fig. 6G, table S6**). Moreover, we observed enrichment for cholesterol metabolism-related genes in Δ*sifA* infected cells (e.g., *Ch25h, Scd2* and *Fabp5*) corresponding to previous reports showing that the loss of vacuolar membrane around SifA mutants requires activity of the cholesterol acyltransferase effector, SseJ (Lossi et al., 2008; Ruiz-Albert et al., 2002). Finally, we studied the effects of Δ*sifA* on host protein levels, using an antibody against Il4ra and flow cytometry. Similar to our results above, we measured high levels of Il4ra protein in Δ*sifA* infected cells only in a subpopulation of cells (**Fig. 6; H and I**). Taken together, using scPAIR-seq we revealed a transcriptional program of host macrophages that is mediated by SifA secreted effector.

## Discussion

Bacterial effectors are key pathogenic virulence determinants that highjack host cellular pathways to support intracellular growth. Despite many efforts, our understanding of the function of many single effector mutants has been limited by their redundancy, heterogeneity, and context-dependent activity. This limitation constrains our ability to discover regulators of bacterial pathogenicity that are the major drivers of infection outcome. In this work we developed scPAIR-seq, in which multiple pooled bacterial mutants can be analyzed simultaneously at single cell resolution to overcome redundancy and heterogeneity during infection. Mutants are functionally characterized by their impact on host responses, that can be expanded to different host contexts. Key to this approach are tagged, multiplexed bacterial barcodes (MIB) that allow us to infect with a pooled library of bacterial mutants and deconvolute mutant identity within single infected host cell. We use this approach to provide a detailed virulence-immune network of single SPI-2 effectors, using a global view of their impact on host gene expression inside infected cells. To overcome expected redundancy between effectors and possible masking of mutant-specific gene signatures by the highly inflammatory gene expression induced upon Toll-like receptor (TLR)-4 detection of bacterial lipopolysaccharide (LPS) (Bode et al., 2012), we developed a computational approach (MAESTRO). Analysis of the impact of each SPI-2 effector on host responses also captures the heterogeneous response of macrophages to single bacterial mutants, exemplified by mutant-specific genes signatures that were expressed in only a subset of infected cells (**Fig. 5; B and C**). The observed variation in the host response highlights the importance of MAESTRO and suggests that T3SS effectors form complex interconnected network that under certain conditions can tolerate perturbations (*e*.*g*. deletion of a single effector) without affecting virulence, as previously suggested (Ruano-Gallego et al., 2021). This redundancy is common to secretion systems of different bacterial pathogen species. We propose that future experiments using scPAIR-seq to specifically analyze cells infected with multiple bacterial mutants or bacteria carrying deletions of multiple effectors can help interpret the combined nonlinear effects of secreted effectors on host processes.

MAESTRO analysis revealed single SPI-2 mutants that impact gene expression of macrophages. We focused on one effector, SifA, a well-studied SPI-2 secreted effector that had most significant impact on host transcriptome in the MAESTRO analysis. SifA has primarily been studied for its role in mediating the formation of a protective niche inside macrophages, and *SifA* deletion results in rupture of the SCV membrane and release of bacteria into the cytosol (Beuzón et al., 2000). However, gene expression changes induced by Δ*SifA* cytosolic *S*.Tm were not studied. Our analysis indicates that cytosolic Δ*SifA S*.Tm drives infected macrophages towards an M2 activation state. Alternatively, vacuolar *S*.Tm with intact SifA, was shown to use another SPI-2 effector to drive M2 activation (Stapels et al., 2018). This is not the case for other cytosolic bacteria, as infection of macrophages with the cytosolic pathogens *Listeria monocytogenes* and *Shigella dysenteriae* was reported to induce an M1-like polarization (Biswas et al., 2007; Lizotte et al., 2014; Rai et al., 2012; Xu et al., 2012). Cytosolic Δ*SifA S*.Tm was shown to expose cytosolic LPS that is detected by Caspase-11 to induce non-canonical inflammasomes and pyroptotic cell death (Aachoui et al., 2013; Thurston et al., 2016). Recently, it was shown that cytosolic LPS can activate phagocytes while retaining their viability (Gaidt et al., 2016), indicating context-dependent cell fate decisions. It is tempting to speculate that in our experimental conditions (low MOI, non-SPI inducing bacterial cultures), macrophages infected with cytosolic Δ*SifA S*.Tm maintain viability and activate a parallel pathway to induce an M2-program. As M2 activation in macrophages is linked to their metabolic state, it would be of interest to further decipher the metabolic pathway that is maintained in SifA infected macrophages.

To conclude, we present scPAIR-seq, an approach to analyze pooled bacterial mutants and cognate host responses at single cell resolution during intracellular infection. At its current scale, scPAIR-seq can be applied for targeted screens of a set of bacterial mutants of interest, as we have done here for SPI-2 *S*.Tm mutants during intracellular infection of macrophages. While SPI-2 mutants have previously been studied, they were mostly analyzed one at a time. Our analysis provides a global view of the functional impact of effectors on immune pathways that underlie intracellular growth and virulence of *S*.Tm. We foresee that enhancing the throughput of scPAIR-seq using microfluidic approaches will allow us to advance our knowledge of fundamental aspects of infection biology at unprecedented level. As discussed above, important points of study will include analysis of the redundancy of bacterial effectors converging on the same immune pathways. Further, experimental models utilizing an intact immune system (*e*.*g*., organoids, blood immune cells or in vivo infections) will provide insight into the impact of bacterial virulence not only on cell autonomous immunity but also their immuno-modulatory capacity in cell-cell communications, and the activity of effectors within different immune cell subsets (Jennings et al., 2017). We propose that at larger scales, scPAIR-seq can be applied as a systematic approach to dissect the interplay between bacterial pathogenesis and host defense strategies in any bacterial strain and infection model.

## Notes

### Competing Interest Statement

The authors have declared no competing interest.

