## Supplementary data for "Paired single-cell host profiling with multiplex-tagged bacterial mutants reveals intracellular virulence-immune networks"

### Supplementary Material & Methods

#### Experimental procedures

##### Bacterial strains and plasmids

*Salmonella enterica* subsp. Typhimurium 14028s (*S.*Tm) wild type (WT) and 24 Single Gene Deletion (SGD) mutants (Porwollik et al., 2014) were used for the MIB-tagged effector mutant library (see table S7) . Deletion was verified by PCR using primers indicated in table S8. To generate MIB-tagged plasmids (pGFP-MIB), a unique MIB sequence of 10 nucleotides followed by a polyA sequence of 32 nucleotides was cloned by restriction-free cloning (van den Ent and Löwe, 2006) downstream to the GFP coding sequence of pFPV25.1 plasmid (Addgene) (see primer list in table S8). Twenty-five pGFP-MIB plasmids were validated and transformed to each of the *S.*Tm 14028 WT and 24 SGD mutants (sequences listed in table S1). We validated no growth impairment for all the pGFP-MIBs strains in liquid culture. For infection of macrophages, *S.*Tm strains harboring the original pFPV25.1 or pGFP-MIBs with a constitutive GFP expression were used for detection of intracellular bacteria. For complementation studies of *SifA*, pSifA was constructed by gene amplification of *sifA* from WT *S.*Tm followed by cloning of the amplified *SifA* gene into pBAD24 (Addgene) using restriction-free cloning. Primers are indicated in table S8.

##### Bacterial growth media

S.Tm strains were grown for 16 hours at 37°C in Luria-Bertani (LB) media and diluted to a similar dilution, based on optical density at 600nm ($OD_{600}$) for further processing. For infection of macrophages, *S.*Tm strains were grown for 14 hours in a SPI-2 inducing media (Stapels et al., 2018): micro oxygen-limited conditions in MgMES media (170 mM 2-(N-morpholino) ethanesulfonic acid (MES) at pH 5.0, 5 mM KCl, 7.5 mM (NH4)2SO4, 0.5 mM K2SO4, 1 mM KH2PO4, 8 mM MgCl2, 38 mM glycerol, and 0.1% casamino acids). 0.2% L-arabinose was added to the growth cultures for induction of *SifA* from pBAD24.

##### GFP fluorescence measurements in MIB-tagged *S.*Tm

LB grown cultures of non-fluorescent *S.*Tm 14028 WT or fluorescent strains carrying either a pFPV25.1 or pGFP-MIB were diluted to $OD_{600}$ of 0.005 and grown in LB in 96-well microplates (Invitrogen M33089) with agitation at 37°C for 24 hours in an imaging multi-mode reader

(Cytation 5). $OD_{600}$ and GFP fluorescence (435/20, 485/20 filter set) were measured every 20 min. In fig. S1A, bacterial fluorescence is presented normalized to $OD_{600}$.

**Quantitative real–time PCR (qRT-PCR) of bacteria expression**

Cultures of *S.*Tm 14028 WT strains carrying either a pFPV25.1 or pGFP-MIB were grown in Luria-Bertani (LB) media for 16 hours at 37°C with shaking, pelleted, washed with phosphate buffer saline (PBS) and resuspended in RLT buffer (Qiagen). RNA was extracted using RNeasy mini kit (Qiagen) with DNase-I treatment (45 min at 23°C) (Qiagen). Resuspended RNA samples were treated with Turbo DNase (30 min at 37°C) (Invitrogen) for complete removal of bacterial DNA. nRNA samples were then processed to capture and amplify only polyA containing transcripts using the CEL-seq2 protocol up to the IVT step (Hashimshony et al., 2016). cDNA was synthesized using Superscript-II (Invitrogen) with random hexamers. Quantification of GFP transcripts was performed by qRT-PCR using GFP primer (table S8) in a QuantStudio 5 instrument (Applied biosystems).

**Bulk RNA-seq analysis for sensitivity of MIB detection**

A mouse J774A.1 macrophage cell line was plated in non-tissue culture treated six–well plates ($4x10^5$ cells/well) supplemented with DMEM containing 10% FBS and 5% sodium pyruvate. Twenty-four hours later, J774A.1 macrophages were infected at MOI of 20:1 with a MIB-*S.*Tm strain grown in LB media at 37°C shaken for 16 hours. Cells were centrifuged to synchronize infection, and 30 minutes later were washed with Gentamicin (Sigma-Aldrich, G1272) to remove non-phagocytosed bacteria. Cells were harvested 4 hours post infection (hpi) by a PBS wash followed by incubation of 10 min with ice-cold EDTA (5mM) and analyzed by flow cytometry (BD FACS Aria III) under continuous cooling at 4°C. Infected (GFP+) cells were sorted into 1.7ml low–bind tubes (Eppendorf) containing 25µl of lysis buffer (0.1% Triton™ X-100 [Sigma] in RNase-free water), placed immediately on dry ice and stored in minus 80.

**Single molecule RNA fluorescence in situ hybridization (smFISH)**

smFISH probes were designed according to Stellaris guidelines, as described in (Raj et al., 2008), synthesized with a 3′ amine modification, covalently labelled with Cy5 mono-reactive dye (GE Healthcare, PA25001), A594 succinimidyl ester (Thermo Scientific, A20004), or TAMRA succinimidyl ester (Thermo Scientific, C6123) as described in (Skinner et al., 2013), and purified through ethanol precipitation. smFISH was performed as described previously (Raj et al., 2006) with few modifications. Briefly, naïve or WT/Δ*sifA* infected cells were cultured in multichambered coverglass, and 20 hpi cells were fixed in 4% PFA (ThermoFisher, 28906), and permeabilized in 70% ethanol at 4°C. Samples were simultaneously co-stained with probes for *Il4ra* mRNA (labelled in TAMRA), *Gapdh* mRNA (labelled in A594), and *Dab2* mRNA (labelled in cy5). Probes were hybridized in humified chambers at 37°C in 15% formamide hybridization buffer (10% dextran sulphate (Sigma D8906), 1 mg/ml E.coli tRNA (Sigma R4251), 2x saline sodium citrate buffer SSC (Ambion AM9765), 2mM vanadyl ribonucleoside complex (NEB S1402S), 0.02% BSA (Thermo Scientific AM2616)) at a final concentration of 300 nM. Before mounting, cells were probed with 1:150 anti-LPS FITC-conjugated antibody (Santa Cruz, sc-52223) for 1h at 37°C, and subsequently washed 3 times for 30 min in wash buffer (2x SSC, 15% formamide), adding 10 mM DAPI (Sigma, D9542) to the third wash to stain the nuclei. Cells were mounted on microscopy slides using GLOX anti-fade buffer (10 mM Tris pH 8, 2x SSC, 0.4% glucose) supplemented with 37 mg/ml glucose oxidase (Sigma, G2133) and 100 mg/ml catalase (Sigma, C3515). Imaging was performed as described below, on the same day as the mounting.

**Microscopy**

For smFISH, an inverted epifluorescence microscope (Eclipse Ti2-E Nikon) equipped with a x100 NA 1.45 oil-immersion objective and a EMCCD camera (iXon Ultra 888, Andor). The filter sets used were 49000-ET (Chroma), 49002-ET (Chroma), 49004-ET (Chroma), 49008-ET (Chroma) and 49006-ET (Chroma) for DAPI, FITC, TAMRA, Alexa 594, and Cy5, respectively. Image stacks consisting of 20 *z* focal planes with 300 nm spacing were acquired for several xy slide positions per sample.

**Image processing for smFISH**

Quantification of RNA FISH images was done using a MATLAB GUI software available at https://github.com/arjunrajlaboratory/rajlabimagetools. Briefly, host cell boundaries are manually identified, and for each fluorescent channel the signal is distinguished from noise through a semi-automated thresholding. Additionally, bacteria coordinates were identified using the LPS signal as input for Schnitzcell (easerver.caltech.edu/wordpress/schnitzcells/) as previously described (Young et al., 2012) and using MATLAB we identified the bacteria within the host cells. Briefly, the bacterial x, y coordinates were extrapolated for each image by finding the centroid of each bacterial object. Host cells were treated as polygons, and the bacterial points were queried to be inside or outside each of the polygon boundaries. The extracted spot counts per host cell, combined with the infection count, were analyzed across samples using R.

**Bone marrow-derived macrophages (BMDMs) isolation**

Bone marrow was extracted from mice as previously described (Falk et al., 1988) Female C57BL/6 mice, aged 6-8 weeks, were euthanized and femurs and tibias were harvested. Bone marrow was flushed from the cells, re-suspended in DMEM, and plated in non-tissue culture treated dishes in DMEM media containing 5% sodium pyruvate, 20% fetal bovine serum, and 25 ng/ml recombinant mouse M-CSF (rmM-CSF, Bio-rad). Cells were harvested at day 6 after bone marrow isolation and were frozen until use. All bone marrow extractions were performed in accordance with the Institutional Animal Care and Use Committee at the Weizmann Institute.

**BMDMs infection with S.Tm for bulk RNA-seq analysis**

BMDMs ($8x10^5$ cells/well) were plated in non-tissue culture treated six–well plates supplemented with DMEM containing 20% FBS and 25 ng/ml rmM-CSF. LB-grown cultures of WT or each of 24 MIB-tagged SPI-2 mutant strains were opsonized with 20% mouse serum (Sigma) for 30 min and washed with PBS. BMDMs were infected with each of SPI-2 mutants separately at MOIs of 5:1 and spun down for 5 min at 400g to synchronize internalization. After 30 min, cells were washed with media containing 50 µg/ml gentamicin to remove *S.*Tm that were not internalized. Fresh media containing 50 µg/ml gentamicin was then added back to the cells for the duration of infection. Infected samples were harvested and sorted by FACS 4 hpi as above. Sorted bulk samples were processed to generate MIB-libraries.

**Bacterial culture for MIB classification**

Twenty-five cultures of the MIB-tagged effectors were grown in LB media at 37°C shaken for 16 hours. Cultures were normalized to the same $OD_{600}$ value and equal volume was taken from all cultures to generate a pooled library of SPI-2 effector mutants. Pelleted pool was re-suspended with RLT buffer (Qiagen), RNA was extracted and MIB-libraries were generated.

**Real-time qPCR of host genes**

BMDMs were infected with WT or Δ*sifA* as described above. At indicated time points, naïve and infected cells were harvested and sorted into 1.7ml low-bind tubes (Eppendorf) containing 700µl of Qiazol® Lysis Reagent, placed immediately on dry ice and stored in -80°C. RNA was extracted using miRNeasy mini kit (QIAGEN, 217004) with DNase-I treatment (QIAGEN). cDNA was synthesized using Superscript-III (Invitrogen). Quantification of transcripts was performed by qRT-PCR using gene-specific primers (table S8) in a QuantStudio 5 instrument (Applied biosystems).

**Antibody staining for flow cytometry**

Infected BMDMs were harvested 20 hpi by a PBS wash followed by 10 min incubation with ice-cold EDTA (5mM), transferred to 1.7ml low–bind tubes (Eppendorf) and spun down for 3 min at 500g, 4°C. Samples were re-suspended with FACS Buffer (PBS, 10 mM EDTA, 1% FBS), transferred to non-treated round bottom 96-well plate (Corning, 3879), washed again with FACS buffer and incubated with 1:50 of CD16/CD32 blocking antibodies (Biolegend, 101301) per sample for 10 minutes on ice. Subsequently, samples were washed with FACS buffer and stained with 1:100 of anti-CD124 (IL-4Rα) PE/Cyanine7-conjugated antibody (Biolegend, 144805) per sample and incubated for 30 minutes on ice. For intracellular LPS staining, samples were fixed in 4% PFA (ThermoFisher, 28906) and permeabilized in BD perm/wash buffer (BD Biosciences, 51-2019KZ) followed by staining with 1:50 anti-LPS FITC-conjugated antibody (Santa Cruz, sc-52223) for 30 min on ice. Samples were then washed with perm/wash buffer, resuspended with FACS buffer, passed through a cell strainer and analyzed by BD FACS Aria III.

**Sample preparation for bulk RNA-seq**

BMDMs were left uninfected (naïve) or infected with WT or Δ*sifA S.*Tm strains for 20h at MOI of 2:1 and samples were sorted and processed to generate host libraries according to the standard Cel-seq protocol (Hashimshony et al., 2016). Libraries were sequenced on an Illumina Novaseq and paired-end sequencing was performed, reading 13 bases for read 1, 6 bases for index 1 and 60 bases for read 2, with a total coverage of ~50M reads per sample.

**scPAIR-seq library preparation for MIB identification in bulk**

scPAIR-seq relies on the CEL-seq library preparation protocol (Hashimshony et al., 2016) up to in-vitro transcription (IVT) step. After IVT, amplified RNA (aRNA) was treated with EXO-SAP enzyme (15 min at 37°C) (Thermo Fisher, 78201) to remove unused primers leftovers. aRNA sample was split and half of the sample volume (11 μl) was used to perform nucleic acid purification with RNAClean XP beads (Beckman Coulter, A63987). The purified aRNA was converted to cDNA using superscript III (Invitrogen, 18080044) with specific GFP primer and cDNA was purified with AMPure XP beads (Beckman Coulter, A63881). MIB library was amplified by 25 PCR cycles using a nested GFP primer to improve specificity of MIB amplification. To ensure exclusive sequencing of MIB reads by a custom GFP primer in the downstream steps, the nested GFP-primer was attached with a 5′-tail containing Illumina 3′-adaptor that was modified in its binding site for the Illumina sequencing primer. Quantification of MIB-library was performed using NEBNext® Library Quant Kit for Illumina® (NEB, E7630L). Libraries were sequenced on an Illumina Miniseq instrument according to standard protocols. Paired-end sequencing was performed with 13 bases for read 1 (R1), 6 bases for index 1 and 60 bases for read 2 (R2). Sequencing was done using custom primer for index 1 to bind the modified adaptor sequence and a custom GFP primer for read 2. All primers are listed in table S8.

**scPAIR-seq library preparation for scRNA-seq experiments**

BMDMs were plated ($8x10^5$ cells/well) and infected at MOI of 2:1 with a pool of opsonized MIB-tagged effector library grown in SPI-2 media at 37°C shaken for 14 hours. At the indicated time points after infection, cells were harvested and sorted either in 1000 cells into 1.7ml low–bind tubes (Eppendorf) containing 25µl of lysis buffer (as described above) for processing of bulk samples or single cells into twin.tec® PCR plate 384 LoBind® (Eppendorf) containing 1.2µl of Cel-seq Lysis buffer (0.2µl of 25ng/µl CEL-seq primer, 0.1µl of dNTPS 10mM [Thermo scientific, # R0181], 0.12µl of 1% Triton™ X-100 [Sigma], 0.012µl of RNnase-in® Plus [Promega, N261B (40U/µl)], 0.768µl Rnase-free water) for single-cell processing. Sorted samples were placed immediately on dry ice and stored in -80°C. To measure bacterial burden, each sorted single-cell was indexed for its GFP intensity levels and analyzed as described in 'GFP indexing data analysis' paragraph. As each 384-well plate was composed of 4 identical sets of 96 Cel-seq primers, single cells were pooled into 4 samples of 96 cells to allow for downstream demultiplexing of libraries by RNA PCR Index Primers (RPIs). Libraries were constructed up to IVT step and samples were split into two parts: 40% of samples volume was used to generate and sequence MIB-libraries as described above. 60% were used to generate host libraries as per the Cel-seq protocol (Hashimshony et al., 2016). Bacterial MIB-libraries were sequenced with a total coverage of ~66M reads from 3648 cells. Host libraries were sequenced on an Illumina Nextseq instrument according to standard protocols. Paired-end sequencing was performed, reading 13 bases for read 1, 6 bases for index 1 and 60 bases for read 2 with a total coverage of ~488M reads from 3264 cells. All the indicated primers are listed in table S8.

#### Bioinformatics analysis

##### Design of multiplex MIB tags

To reduce ambiguity of MIB identification due to sequencing errors, we selected barcodes that allow for at least 3 mismatches. The string distance between all barcodes was computed as the generalized Levenshtein (edit) distance, giving the minimal number of insertions, deletions and substitutions needed to transform one string into another. The edit distance between all pairs of bacterial barcodes was designed to be larger than 2.

##### Processing of bacterial MIB reads

Fastq reads were demultiplexed to samples (bulk libraries) or cells (single cell libraries) based on the CEL-Seq barcode located in positions 7-12 of R1. R2 reads were filtered to contain the GFP sequence preceding the MIB (positions 1-37 of R2; GFP sequence: TGGGATTACACATGGCATGGATGAACTATACAAATAA, for Δ*sseJ* the sequence is TGGGATCACACATGGCATGGATGAACTATACAAATAA due to mutation in the plasmid during cloning). To account for sequencing errors, 1 mismatch was allowed for the GFP sequence identification. The sequence of 10 nucleotides following the GFP (positions 38-47 of R2) is the bacterial barcode, i.e., the MIB sequence. In each sample/single-cell, we counted the number of reads that correspond to each bacterial barcode allowing 1 mismatch in the MIB sequence (see the specificity section below). For bulk libraries we collapsed read count to UMI count to avoid biases due to PCR amplification. Using this procedure, we generated a matrix with the counts of each mutant (MIB) in each cell/sample.

##### Specificity of MIB classification

To evaluate the specificity of correct MIB identification we iteratively increased the number of mismatches that are allowed in order to associate a read to a specific mutant (MIB). We calculated the number of reads that are assigned to each mutant with perfect match (0 mismatches) and up to 10 mismatches. The curve of each mutant (fig. S2C) is in agreement with the edit distance of the MIB to all other mutants. In addition, this analysis validates that allowing 1 mismatch in the MIB classification increases the number of detected reads without affecting robustness and specificity of correct MIB identification.

**Noise estimation for MIB classification**

For MIB-library generation infected cells/samples are pooled. During the PCR amplification artifacts can be formed due to template switching between homologous transcripts (Kanagawa, 2003). Using bulk RNA-seq data from BMDMs infected with each SPI-2 mutant alone, we evaluated this noise between mutants. Based on the MIB matrix generated as described above in 'Processing of bacterial MIB reads', we counted the percentage of false positives for each mutant (fig. S2D). Comparing the correct MIB identification to the false positive rates validated specific MIB identification for all infected mutants.

**Removal of ambiguous bacterial reads in single cell libraries**

Since each 384-plate was divided to 4 pooled samples of 96 single cells with identical CEL-seq primers (as described above), different RPIs were used for each pool to enable demultiplexing of single cells. During demultiplexing, index hopping can lead to assignment of sequencing reads to the wrong index in low percentage (Kircher et al., 2012). To eliminate MIB identification due to wrong assignment of reads to the correct index, we filtered all reads with identical CEL-Seq barcode, UMI and bacterial barcode (perfect match of all fields) that originated from 2 or more different plates in the same sequencing run. We included in the analysis only reads that originate from the plate with the highest count of these reads, to eliminate ambiguous reads. In case of equal number of ambiguous reads from the different plates, all duplicate reads were removed. In addition, reads with one or more 'N' nucleotide in their UMI sequence (positions 1-6 of R1) were removed to avoid ambiguous read assignment.

**Mutant detection from bacterial single cell libraries**

To identify the identity of mutants in each single infected cell, we performed a global analysis that is based on the distribution of all MIBs across all single cells. Cells with less than 128 MIB reads in total were removed from the analysis due to low coverage (fig. S3A; 1,191 cells remain). MIB matrix was normalized by 'NormalizeData' function from 'Seurat' (Hao et al., 2021) using the CLR method (centered log ratio transformation), and batch effect between plates was removed by 'removeBatchEffect' function from 'limma' library (Ritchie et al., 2015). Then, to detect which mutant infected each cell, we applied 'deMULTIplex' R package (McGinnis et al., 2019). Using this procedure we modeled the probability density function of each normalized MIB distribution across all cells, and found the local maximum of the positive and negative cells (highest and lowest maximum, respectively) for each MIB alone. To set a threshold for positive and negative cells for each MIB we optimized the quantile parameter between the maxima of each MIB which maximizes singlet detection across all cells in the experiment. Using this optimized quantile, we assigned each cell to the mutant (or mutants) it was infected with.

**Mutant detection from host libraries**

As a validation for mutant detection from MIB libraries, we searched for GFP reads in host libraries. The GFP sequence can be found in any position of R2 read due to the fragmentation step in the standard CEL-Seq protocol of the host samples. Fastq reads were filtered to contain the GFP sequence TGGATGAACTATACAAATAA, and the following 10bp after the GFP sequence were considered as the MIB sequence. Cells with 1 or more GFP reads that their MIB sequence matched a specific bacterial barcode with up to 1 mismatch were considered as infected with the corresponding mutant. Cells with more than 1 detected mutant were considered as infected with multiple mutants. The identification of MIB based on host library was used only to validate the bacterial libraries and assess its improved detection efficacy, and not for downstream analysis.

**GFP indexing data analysis**

We used the 'flowCore' R package to extract the GFP intensity of each single cell from the flow cytometry data (FCS files).

**Host single cell RNA-Seq data preprocessing and normalization**

The CEL-Seq pipeline (https://github.com/yanailab/CEL-Seq-pipeline) was used for single cells demultiplexing, alignment to mm10 genome and UMI counts. Overall we sequences 3264 cells, with 136,173 mean reads per cells, 14,929 median UMI count per cell and 3835 median genes per cell. Cells with low coverage (less than 3000 UMIs) and high percent of mitochondrial genes (more than 10%) were removed from downstream analysis. Data was normalized to cell size factors. Size factors were calculated using 'computeSumFactors' function of R library 'scran' (L. Lun et al., 2016) and data normalization was performed using the 'normalize' function of R library 'scater' (McCarthy et al., 2017). Data was log2 transformed with pseudo-count of 1.

**Host single cell RNA-seq data analysis**

The analysis was done only on the host transcriptome of the 850 single cells for which we identified the invading mutant (using our pipeline) and passed quality control of the host single cell data. We filtered out cell cycle and ribosomal genes, and selected the top variable genes (523 genes) based on the mean expression and dispersion of genes (coefficient of variance) across all single cells. Principal component analysis (PCA) was performed on the variable genes, and the first 26 PCs were selected for downstream analysis based on the inflection point of the curve of the variance explained by each successive principal component. Unsupervised clustering of the single cells was done using Louvain community detection (Blondel et al., 2008) on the k-nearest neighbor (KNN) graph, which was generated with k=20 on the Euclidian distance between cells in the PC space. We obtained 6 clusters that were not equally distributed across the 24 mutants and WT infected cells.

**Host single cell transcriptome stability analysis**

To evaluate the number of cells required from each SPI-2 mutant in order to detect robust changes in host transcriptome, we modeled the variance across genes for increasing number of infected single cells. We generated 1000 random groups with 2 to 200 cells, calculated for each group the mean variance of all genes, and tested how many cells are required to stabilize the gene variance. According to this analysis the variance is starting to stabilize around 20 cells, fitting with our mutant groups which contain 19-76 infected cells per group.

**MAESTRO analysis**

To identify the differences between host response of cells infected with WT *S*.Tm versus SPI-2 individual mutants we performed supervised analysis on the host transcriptome which compares WT infected cells to each SPI-2 mutant. Cells that were infected with the same invading mutant were grouped (24 mutant groups and a WT group) and mean expression and coefficient of variance (CV) were calculated for each gene in each group, on the log2 transformed data. To compare between WT and specific mutant infected cells we developed a metric called MAESTRO (MutAnt spEcific hoSt TRanscriptOme) which captures two important aspects of gene expression in single cell data: *1)* change in expression levels between cells (i.e., difference in mean expression between WT and specific mutant infected cells) *2)* change in expression variability across cells (difference in CV between WT and specific mutant infected cells). The statistic of our method is the MAESTRO score:

$$MAESTRO_{gene\ i} = \sqrt{w(X_{WT} - X_{Mut\ j})^2 + (Y_{WT} - Y_{Mut\ j})^2}$$

Where $X$ is the mean expression of *gene i* in the WT ($X_{WT}$) or specific mutant ($X_{Mut\ j}$) infected cells, and $Y$ is the CV of *gene i* across the WT ($Y_{WT}$) or specific mutant ($Y_{Mut\ j}$) infected cells. $w$ is a weight for the mean expression parameter in order to control for artificial high CV that results from lowly expressed genes due to dropouts (w=1.5).

To further exclude noise coming from lowly detected genes we calculated the MAESTRO score only for genes that are expressed with at least 2 UMIs from at least 30% of the cells (WT group or in a specific mutant). If the gene is expressed only from one group, the mean and CV values in the missing group are set to the minimal values.

The distribution of MAESTRO scores across all expressed genes from all mutants generated a right tail of genes with high scores. We defined the 95$^{th}$ percentile of the distribution as the threshold to define a gene as differentially expressed between specific mutant and WT infected cells. The list of differentially expressed genes (DEGs) for each mutant relative to the WT was generated using this threshold. The number of DEGs for each mutant varied from 27-240 genes. 32 genes were shared between the DEGs lists of 10 mutants or more (11 up and 21 down-regulated genes). We excluded these genes from all lists since we suspected that these genes have artificial high MAESTRO scores.

To provide statistical power and test for mutants that significantly differentiate host transcriptome relative to WT we generated a random model. We included in our random model all cells that were excluded before due to low MIB coverage (less than 128 total MIB count), and therefore the identity of the infecting mutant was unknown (2457 cells). We assigned these cells randomly to 25 groups in the same size as our original data (fig. S3C) and calculated the MAESTRO score for all genes in all mutants' groups. We repeated this procedure 1000 times to generate the null distribution for each mutant. Then, two-sample t-test was performed to compare between the MAESTRO scores of each mutant to its null distribution. Overall, 6 mutants significantly elevated the MAESTRO scores distribution relative to their null distribution (1%FDR), and thus significantly affected host transcriptome relative to WT infected cells: Δ*sifA*, Δ*gogB*, Δ*sopD2*, Δ*ssaV*, Δ*sseK1* and Δ*srfJ*.

**Proportion of host cells expressing a specific signature**

To classify whether a signature is up or down regulated in specific single cell, we calculated the mean expression of the genes included in the signature across all cells. Next, we transformed mean expression levels to z-scores and set a threshold to define cells that elevate the signature (z-score>1.5) or reduce the signature (z-score<-1.5).

**Bulk RNA-seq data analysis**

The CEL-seq pipeline (https://github.com/yanailab/CEL-Seq-pipeline) was used for sample demultiplexing, alignment to the genome (mm10) and gene counting. Each condition had 4 replicates; one replicate of naïve cells was excluded due to low coverage (~2M reads). The package DESeq2 (Love et al., 2014) was used to normalized the data and identify differentially expressed genes between naïve, WT and Δ*sifA* infected cells. Lowly expressed genes (mean expression across all conditions < 3 in log2 scale) were excluded from the analysis.

#### Supplementary figures legend

**Figure S1: Polyadenylation enables specific MIB amplification in *S*.Tm. (A)** GFP fluorescence levels (normalized to $OD_{600}$) of indicated *S*.Tm strains grown in Luria-Bertani (LB) in 96-well plates, with shaking, at 37°C. Lines represent the mean and SEM of 8 replicates, values were measured every 20 min. **(B)** Cultures of *S*.Tm control or MIB-*S*.Tm were harvested for RNA extraction and capture of polyA-containing transcripts. GFP transcripts were detected by specific primers and analyzed by qRT-PCR.

**Figure S2: Validating sensitivity and accuracy of MIB detection. (A)** J774.1 macrophages were infected for 4 hours with MIB-*S*.Tm at MOI of 20:1 and harvested for RNA-seq. After the IVT step, bacterial MIB-libraries were generated from serial dilutions of amplified RNA (aRNA) and proportion of MIB reads from total reads was quantified for each library. **(B)** Edit distance between all pairs of multiplexed MIBs used for SPI-2 mutant library. The edit distance is calculated as the minimal number of insertions, deletions or substitutions needed to transform one MIB into another (see colorbar to the right). **(C)** Cultures of multiplexed-tagged SPI-2 mutants were pooled and MIB library was generated. MIB sequences were extracted from each read and assigned to the matched bacterial barcode with increasing number of allowed mismatches. Presented are the number of reads that were classified to each mutant from 0 to 10 mismatches. **(D)** BMDMs infected with individual SPI-2 mutant were sorted separately and analyzed by MIB libraries. For each SPI-2 mutant we calculated the number of true positives (correct MIB identification) and false positives classifications. To estimate false positive rates without MIB, BMDMs were infected with *S*.Tm control. Presented are the percentage of correct MIB identification for each mutant, and the rates of false positive classification for each mutant, from total GFP reads. For all mutants we have high true positive rates relative to false positive identification.

**Figure S3: MIB identification within single infected cells. (A)** Distribution of total MIB counts ($\log_2$) across all single cell (n=3648). Cells with low MIB coverage were excluded from further analysis (2457 cells; dashed black line). **(B)** Measurement of bacterial GFP indexing, indicated by boxplots that represent GFP intensity levels, as detected by flow cytometry, at 4 hpi. (n=2583) and 20 hpi (n=898). The box represents the median and 25-75$^{th}$ percentile, whiskers encompass all data points, and outlier marked as dots **(C)** Number of single cells that are infected with WT or the different SPI-2 mutants, as identified by scPAIR-seq analysis. **(D)** Comparison of MIB detection by host and bacterial libraries. Most invading mutants were identified only in bacterial library (80%; blue), 19% were identified also by host library and were similar to the identified mutant in the MIB libraries (turquoise), and only 1% of identified mutants were different by host and MIB libraries (yellow).

**Figure S4: Unsupervised single cell analysis reveals both redundancy and mutant-specific fingerprints. (A-C)** Quality control of the host single cell RNA-seq data. Presented are boxplots of the number of genes (**A**), sum of UMIs (**B**) and proportion of mitochondrial genes (**C**) per cell. On each box, the central mark indicates the median, and the box the 25th-75th percentile; whiskers encompass all data points and outlier marked with +. **(D)** Sorted expression matrix of 100 marker genes for the 6 clusters obtained using Louvain community detection on the host single cell data (colorbar to the right indicates relative expression levels). The cells are ordered by their cluster (see colorbar at the bottom). **(E)** Clusters composition for each group of cells infected by WT *S*.Tm or SPI-2 mutants. Presented are proportion of cells from each cluster in each SPI-2 mutant group and WT infected cells. **(F)** To estimate the number of required cells for each SPI-2 mutant in order to detect robust changes in host transcriptome, we performed stability analysis (see supplementary methods). Boxplots represent the variance between genes in random groups of infected cells from 2 to 200 cells. For each box there are 1000 random assignments of cells. Inner plot: zoom-in to groups of 2 to 50 cells showing that from 20 cells gene variance across cells is stabilized.

**Figure S5: Differentially expressed genes between WT and SPI-2 mutant infected cells as detected by MAESTRO analysis.** For each mutant, presented is the mean expression (x-axis) and CV (y-axis) of all expressed genes across all cells infected by this mutant (gray dots). Genes with MAESTRO score above threshold (higher than the $95^{th}$ percentile of all MAESTRO scores across all mutants; see Fig. 4B) are highlighted for each mutant (colored dots). These genes were classified as the DEGs of each mutant.

**Figure S6: Δ*sifA*-infected cells are polarized into M2-transcriptional state at later infection stages:** **(A)** Microscope images of smFISH of Gapdh (left panels), Il4ra (middle panels) and Dab2 (right panels) mRNAs in WT (upper row) or Δ*sifA* infected cells (lower row). Each mRNA is shown in black in the single channel image, or in white overlaid with DAPI to mark the nuclei (in blue), and with LPS to mark intracellular bacteria (in red). Scale bar: 5 μm **(B)** Violin plots showing Gapdh mRNA counts per host cell for each infected sample. The median is indicated by the black diamond **(C)** Il4ra (left panels) and Dab2 (right panels) mRNA counts per host cell infected with WT (upper row) or ΔsifA (lower row). The cells were grouped according to how many bacteria they were infected with (6 groups ranging: 0, 1, 2, 3-5, 6-10, 11-21 intracellular bacteria). The box represents the median and 25-75th percentile **(D and E)** BMDMs were infected with WT or Δ*sifA* at MOI of 2:1 and cells were collected at indicated time point for qRT-PCR analysis with primers specific to M1 **(D)** and M2 **(E)** genes and expression was compared to t=0 (naïve). Lines represent the mean fold change from naïve of 3 replicates and presented also are the standard error (SEM) for the replicates. Expression analysis was normalized to *Rps*13 gene.

### Figure S1

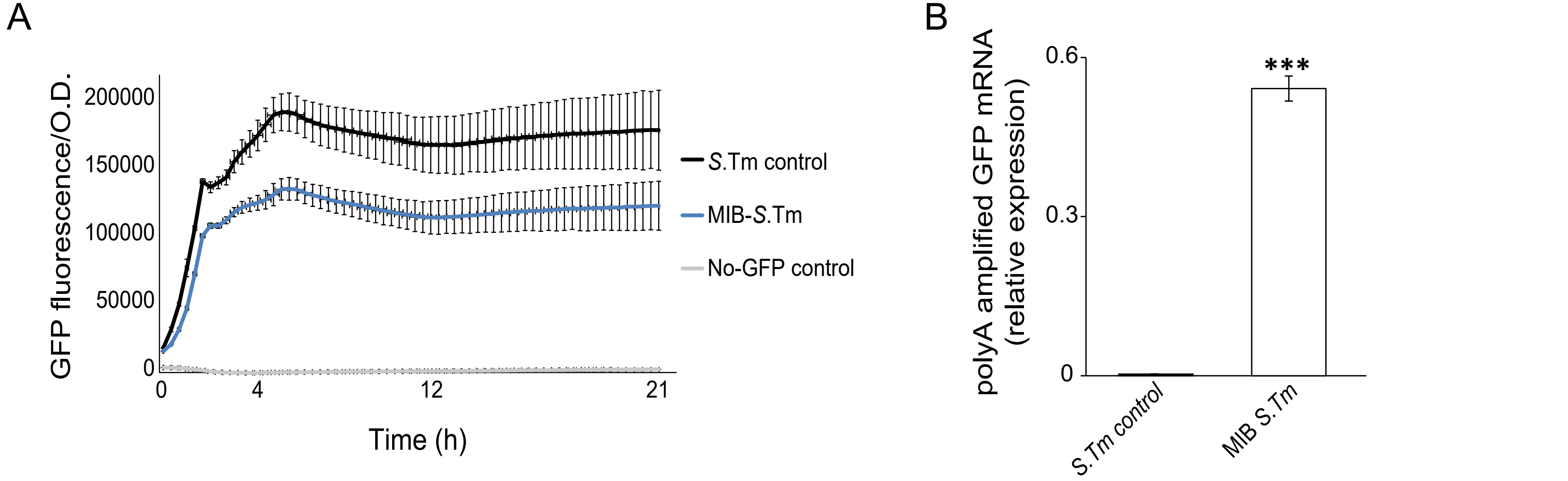
A
GFP fluorescence/O.D.
200000
150000
100000
50000
0
0
4
12
21
Time (h)
S.Tm control
MIB-S.Tm
No-GFP control
B
polyA amplified GFP mRNA
(relative expression)
0.6
0.3
0
***
S.Tm control
MIB S.Tm

Figure S2

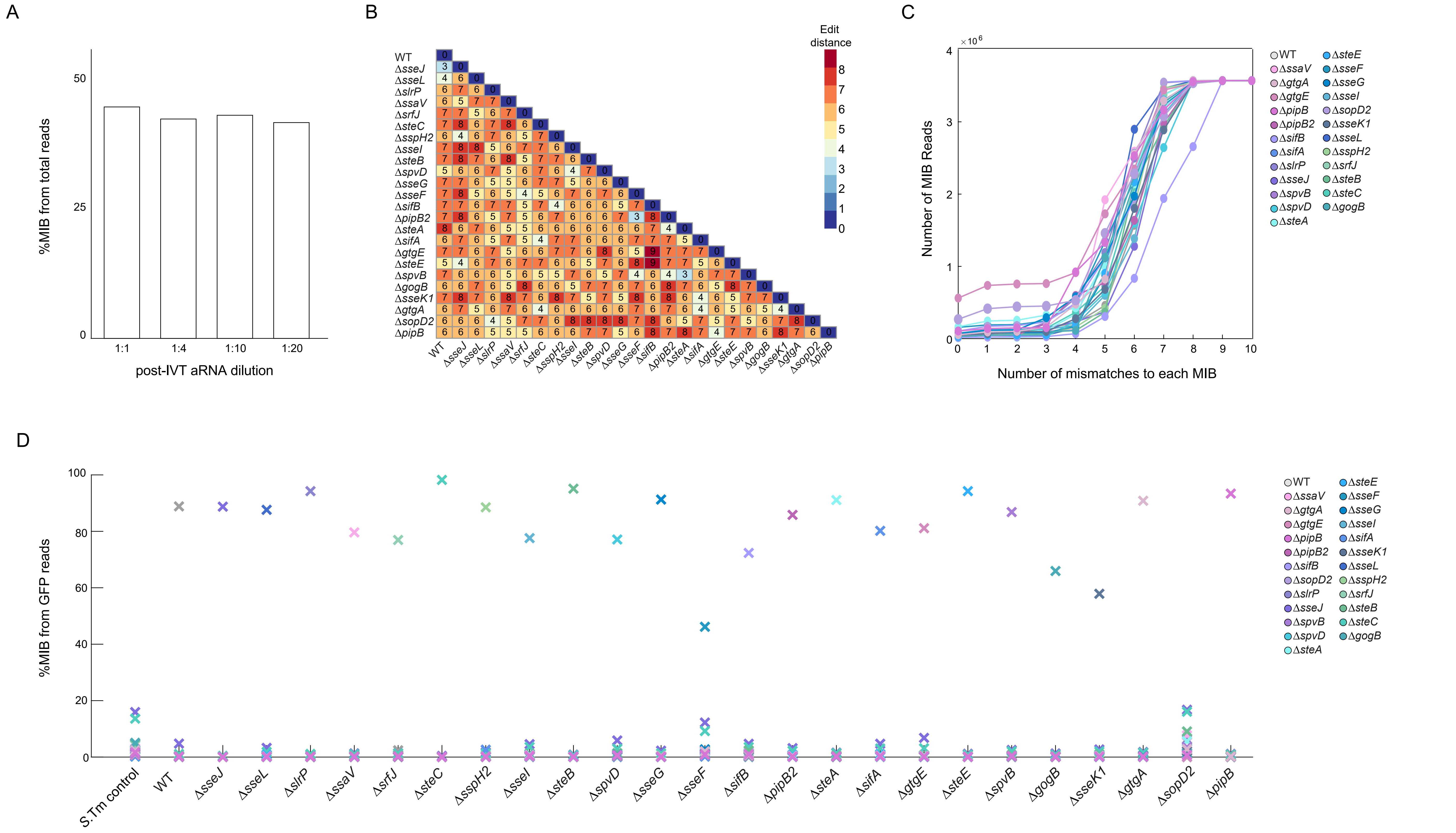
A
%MIB from total reads
50
25
0
1:1
1:4
1:10
1:20
post-IVT aRNA dilution
B
Edit distance
WT
ΔsseJ
ΔsseL
ΔslrP
ΔssaV
ΔsrfJ
ΔsteC
ΔsspH2
ΔsseI
ΔsteB
ΔspvD
ΔsseG
ΔsseF
ΔsifB
ΔpipB2
ΔsteA
ΔsifA
ΔgtgE
ΔsteE
ΔspvB
ΔgogB
ΔsseK1
ΔgtgA
ΔsopD2
ΔpipB
C
Number of MIB Reads
× 10^6
Number of mismatches to each MIB
D
%MIB from GFP reads
S.Tm control
WT
ΔsseJ
ΔsseL
ΔslrP
ΔssaV
ΔsrfJ
ΔsteC
ΔsspH2
ΔsseI
ΔsteB
ΔspvD
ΔsseG
ΔsseF
ΔsifB
ΔpipB2
ΔsteA
ΔsifA
ΔgtgE
ΔsteE
ΔspvB
ΔgogB
ΔsseK1
ΔgtgA
ΔsopD2
ΔpipB
WT ΔsteE
ΔssaV ΔsseF
ΔgtgA ΔsseG
ΔgtgE ΔsseI
ΔpipB ΔsifA
ΔpipB2 ΔsseK1
ΔsifB ΔsseL
ΔsopD2 ΔsspH2
ΔslrP ΔsrfJ
ΔsseJ ΔsteB
ΔspvB ΔsteC
ΔspvD ΔgogB
ΔsteA

### Figure S3

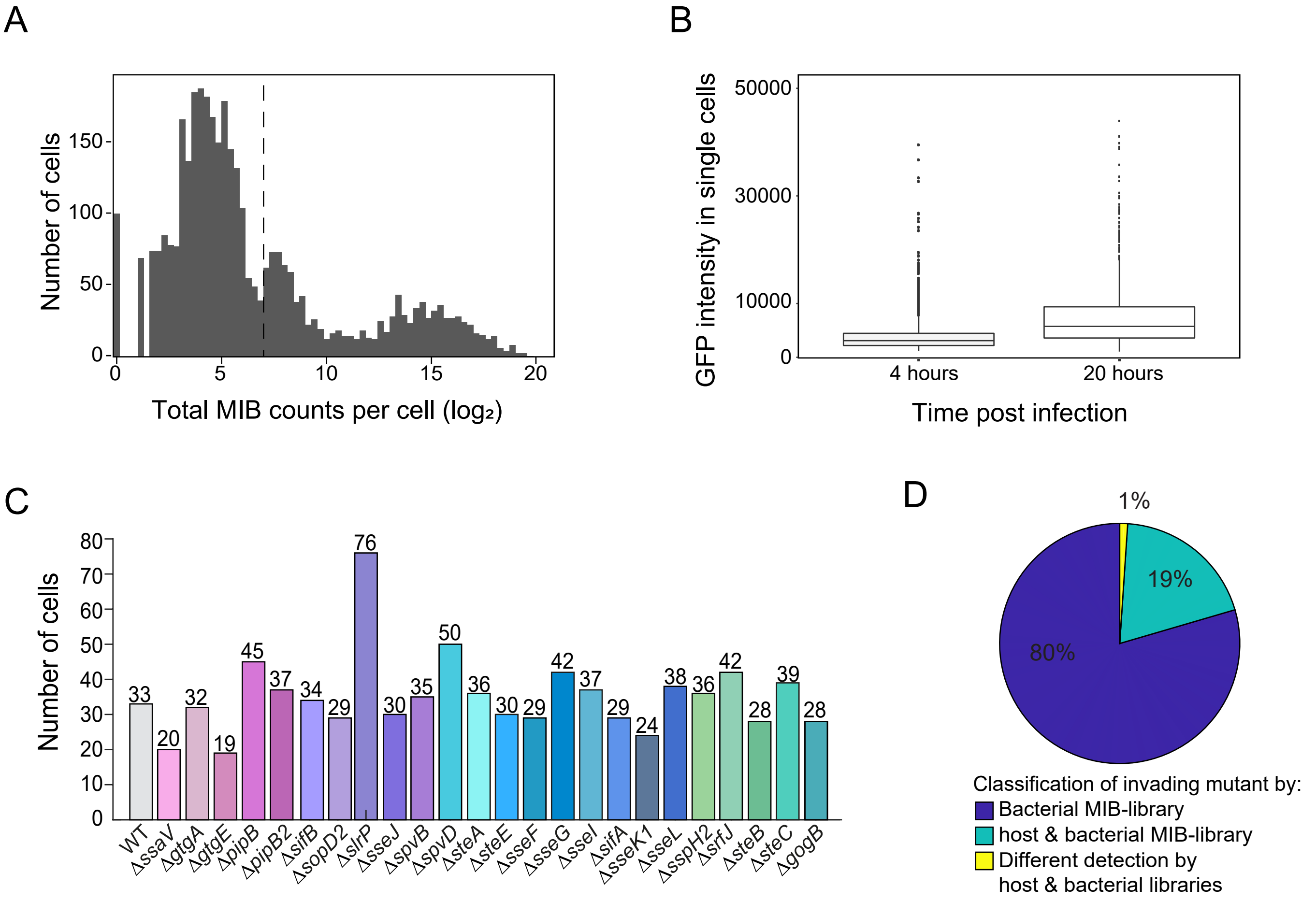

A
B
C
D
Number of cells
150
100
50
0
0
5
10
15
20
Total MIB counts per cell (log₂)
GFP intensity in single cells
50000
30000
10000
0
4 hours
20 hours
Time post infection
80
70
60
50
40
30
20
10
33
20
32
19
45
37
34
29
76
30
35
50
36
30
29
42
37
29
24
38
36
42
28
39
28
WT
ΔssaV
ΔgtgA
ΔgtgE
ΔpipB
ΔpipB2
ΔsifB
ΔsopD2
ΔslrP
ΔsseJ
ΔspvB
ΔspvD
ΔsteA
ΔsteE
ΔsseF
ΔsseG
ΔsseI
ΔsifA
ΔsseK1
ΔsseL
ΔsspH2
ΔsrfJ
ΔsteB
ΔsteC
ΔgogB
1%
19%
80%
Classification of invading mutant by:
Bacterial MIB-library
host & bacterial MIB-library
Different detection by
host & bacterial libraries

Figure S4

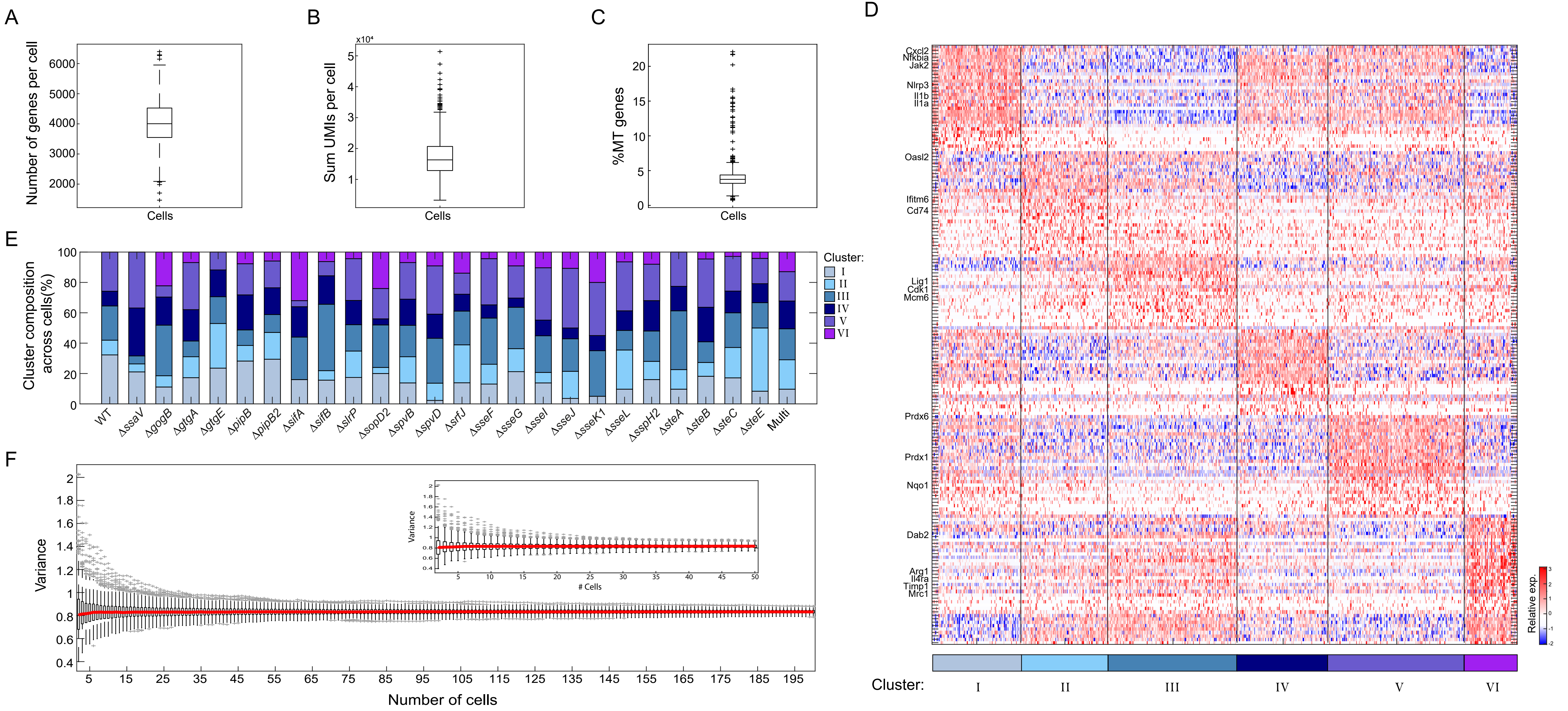
A
Number of genes per cell
Cells
B
Sum UMIs per cell
Cells
C
%MT genes
Cells
D
Cxcl2
Nfkbia
Jak2
Nlrp3
Il1b
Il1a
Oasl2
Ifitm6
Cd74
Lig1
Cdk1
Mcm6
Prdx6
Prdx1
Nqo1
Dab2
Arg1
Il4ra
Timp1
Mrc1
Relative exp.
Cluster:
I
II
III
IV
V
VI
E
Cluster composition across cells(%)
Cluster:
I
II
III
IV
V
VI
WT
ΔssaV
ΔgogB
ΔgtgA
ΔgtgE
ΔpipB
ΔpipB2
ΔsifA
ΔsifB
ΔslrP
ΔsopD2
ΔspvB
ΔspvD
ΔsrfJ
ΔsseF
ΔsseG
ΔsseI
ΔsseJ
ΔsseK1
ΔsseL
ΔsspH2
ΔsteA
ΔsteB
ΔsteC
ΔsteE
Multi
F
Variance
Number of cells
### Cells

### Figure S5

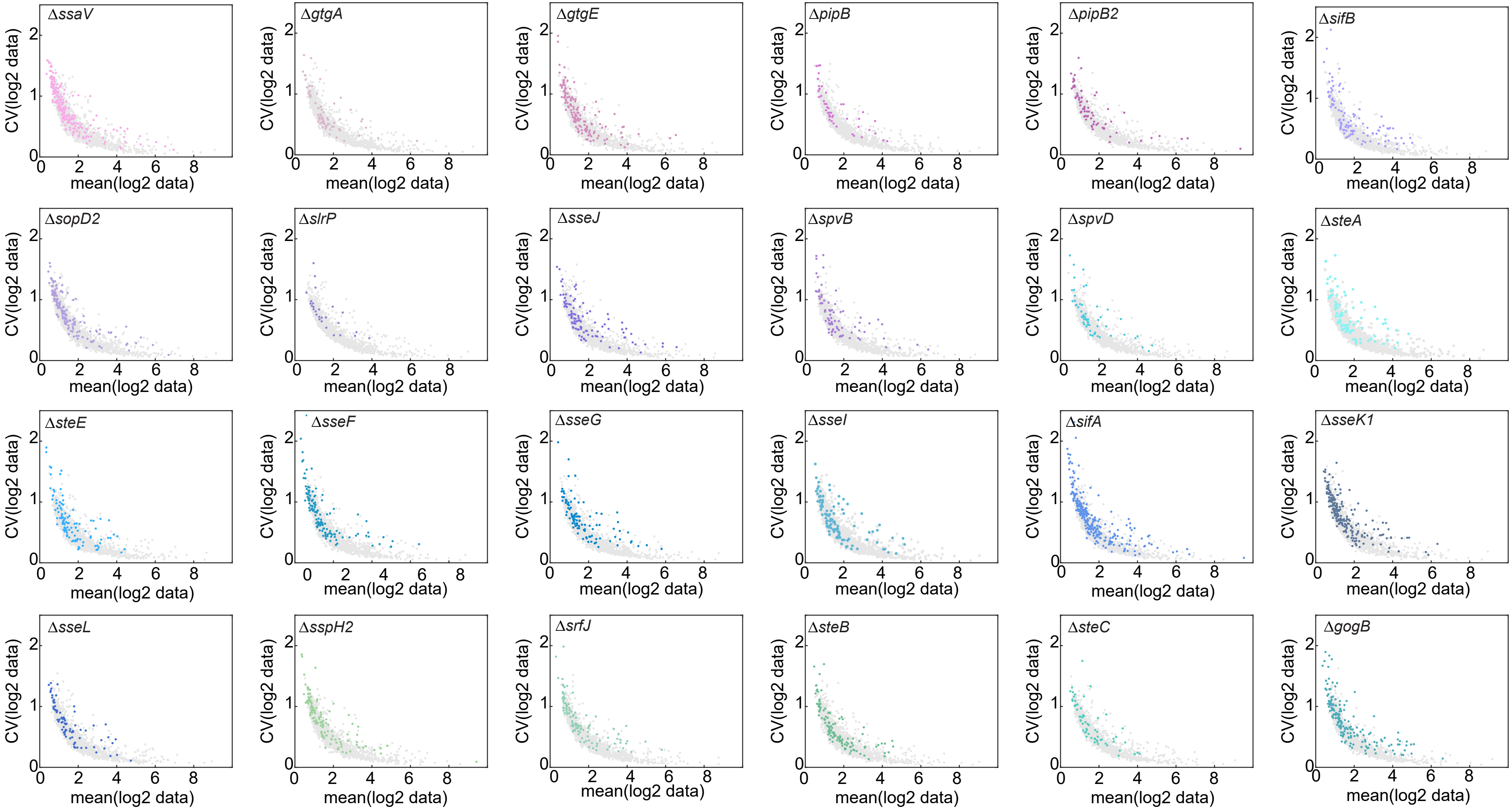
ΔssaV
ΔgtgA
ΔgtgE
ΔpipB
ΔpipB2
ΔsifB
ΔsopD2
ΔslrP
ΔsseJ
ΔspvB
ΔspvD
ΔsteA
ΔsteE
ΔsseF
ΔsseG
ΔsseI
ΔsifA
ΔsseK1
ΔsseL
ΔsspH2
ΔsrfJ
ΔsteB
ΔsteC
ΔgogB
CV(log2 data)
mean(log2 data)

Figure S6

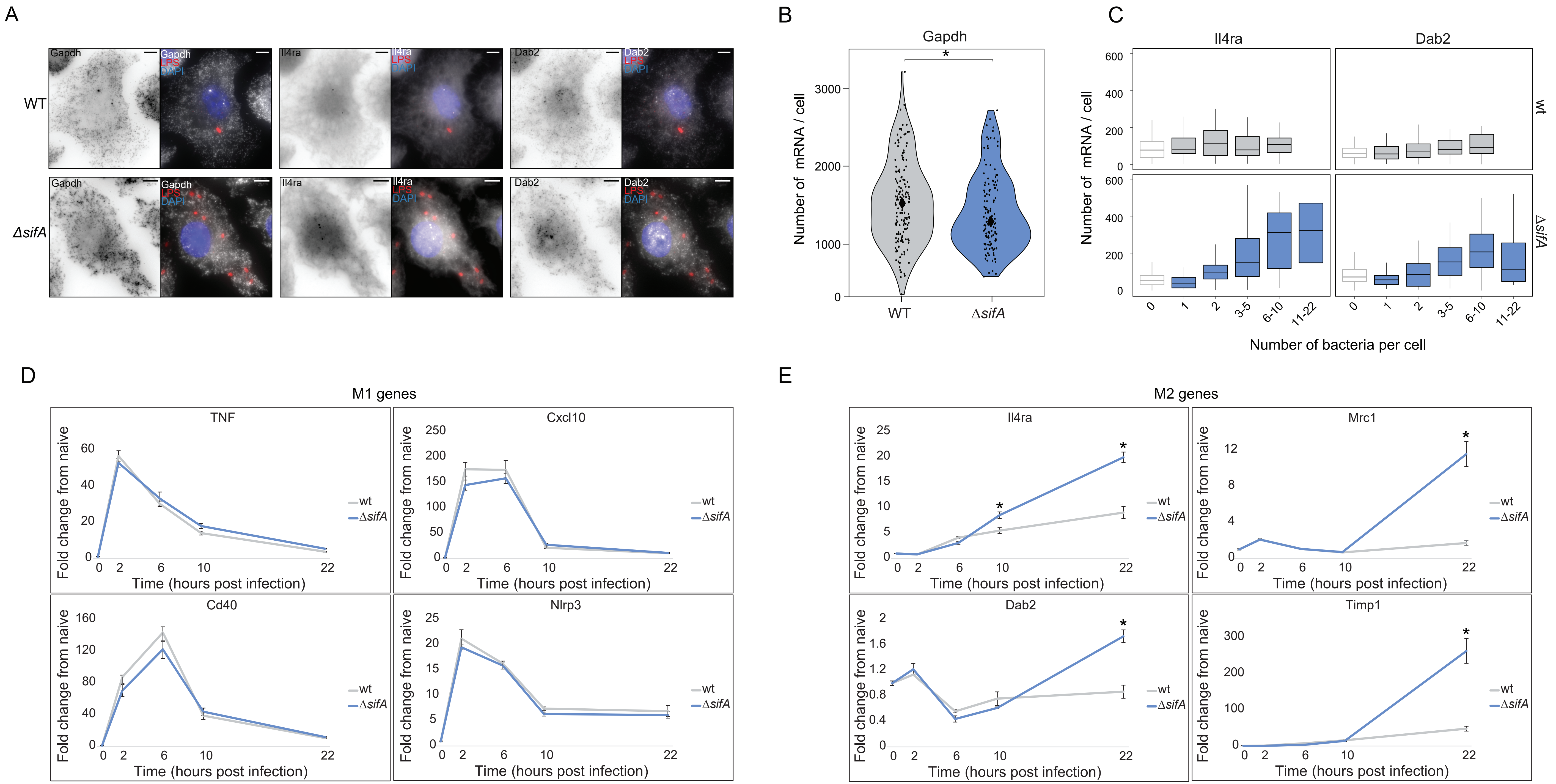
A
B
C
D
E
WT
ΔsifA
Gapdh
LPS
DAPI
Il4ra
Dab2
Number of mRNA / cell
Number of bacteria per cell
wt
M1 genes
M2 genes
TNF
Cxcl10
Cd40
Nlrp3
Mrc1
Timp1
Fold change from naive
Time (hours post infection)
ΔsifA
